# A SET domain-containing protein and HCF-1 maintain transgenerational epigenetic memory

**DOI:** 10.1101/2025.03.25.645221

**Authors:** Chenming Zeng, Giulia Furlan, Miguel Vasconcelos Almeida, Juan C. Rueda-Silva, Jonathan Price, Jonas Mars, Pedro Rebelo-Guiomar, Meng Huang, Shouhong Guang, Falk Butter, Eric A. Miska

## Abstract

Epigenetic information can be inherited through a process known as transgenerational epigenetic inheritance (TEI). TEI can be initiated and maintained by small RNAs and histone modifications. The latter includes histone methylation, deposited by proteins with methyltransferase activity, including SET domain-containing proteins. Other SET domain-containing proteins with no catalytic methyltransferase activity also adopt roles in chromatin and gene expression regulation. Here, we describe SET-24, a SET domain-containing protein in *Caenorhabditis elegans* that belongs to a family of catalytically inactive SET proteins. SET-24 localises to germline nuclei and is required for germline immortality. Additionally, the inheritance of small RNA-driven epigenetic silencing is compromised in *set-24* mutants. Using quantitative proteomics and yeast two-hybrid assays, we found that SET-24 interacts with Host Cell Factor 1 (HCF-1), a protein involved in epigenetic regulation and associated with known chromatin remodelling complexes, like COMPASS, which deposits H3K4me3. In *set-24* mutants, hundreds of genes display increased H3K4me3 levels at their transcription start sites. While these changes are not matched at the transcriptional level, small RNA production is disrupted in approximately one fifth of those genes, which are normally targeted by small RNA pathways. We propose that SET-24 is a factor required to maintain epigenetic memory in the germline by maintaining a chromatin environment permissible to small RNA biogenesis over generations.

## Introduction

RNA interference (RNAi) is a highly conserved gene silencing mechanism present in a wide array of organisms, safeguarding genome integrity and modulating gene expression^1,2^. This process can be triggered by both exogenous small interfering RNAs (exo-siRNAs) and endogenous small RNAs, including endo-siRNAs and Piwi-interacting RNAs (piRNAs). In the nematode *Caenorhabditis elegans*, RNAi involves the synthesis of primary siRNAs, which cleave their target mRNAs within the cytoplasm. This initiates the amplification of secondary siRNAs, also known as 22G-RNAs, bound to Argonaute proteins, thereby amplifying the signal and leading to post-transcriptional gene silencing^3^. Additionally, within the nucleus, small RNAs and Argonaute proteins can promote the deposition of histone modifications such as H3K9me3 and H3K27me3, resulting in transcriptional gene silencing of target genes^4–8^.

Both exogenous and endogenous RNAi cause heritable gene silencing. For example, the silencing effects initiated by GFP-targeting double-stranded RNAs (dsRNAs) can last for multiple generations^9,10^. Argonaute-bound siRNAs and histone modifications such as H3K9me3, H3K23me3, and H3K27me3 have been implicated in the transmission of silencing across generations, a phenomenon also known as transgenerational epigenetic inheritance (TEI)^4,6,7,10–16^. Numerous factors related to histone modifications, such as H3K27me3 demethylation factors, the PRC2 complex, the putative H3K9 methyltransferases MET-2, SET-25, and SET-32, as well as SET-21, which methylates H3K23 in conjunction with SET-32, are involved in TEI^12,17–24^.

In transgenic *C. elegans* with a *gfp::h2b* sequence controlled by a germline-specific promoter, exposure to *gfp* dsRNA results in the silencing of GFP, which is typically inherited for multiple generations^9^. Previous research has proposed three distinct steps to small RNA-driven TEI: initiation, establishment, and maintenance^18^. The establishment of TEI relies on essential chromatin modifiers, specifically SET-25 and SET-32^18^. Other factors, such as HRDE-1, are involved in both the establishment and maintenance of TEI^25–27^.

The mortal germline (Mrt) phenotype in *C. elegans* is a heritable trait characterized by progressive sterility across successive generations^28,29^. Over the past decades, numerous Mrt mutants have been identified, many of which are associated with RNAi and TEI^27^. For instance, mutations in the WAGO-4/ZNFX-1 complex, SET-25, and SET-32, and some nuclear RNAi pathway factors such as HRDE-1, lead to Mrt phenotype in *C. elegans* grown at 25°C^17–19,25,30–32^. It has been hypothesized that RNAi machinery might promote germline immortality by establishing an epigenome conducive to germ cell quiescence^30^. However, the underlying mechanisms for defects in nuclear RNAi or TEI, simultaneous to temperature-sensitive Mrt phenotype, remain unclear. For piRNA-driven silencing, aberrant silencing of histone genes in piRNA mutants was proposed to underlie their Mrt defect^33^.

Wild isolates of *C. elegans* offer a rich source of natural genetic variation for investigating worm behaviours, phenotypes, and their underlying mechanisms. Some wild isolates exhibit a temperature-sensitive Mrt phenotype at 25°C^34^. In a previous study, a deletion of the *set-24* gene was identified in the wild *C. elegans* isolate MY10, which displayed a Mrt phenotype^35^. SET-24 encodes a protein that contains a conserved SET domain, originally identified from Su(var)3-9, Enhancer of zeste, and Trithorax histone methyltransferases (HMTs)^36^. SET domain-containing proteins are a well-characterized group of histone-modifying factors that play a critical role in gene regulation processes. Dysfunction of these factors is frequently associated with diseases, including cancer, developmental disorders, and aging-related pathologies^37^. Unlike the well-established functions of *C. elegans* SET-25 and SET-32 in H3K9 methylation and TEI, the function and mechanism of SET-24 remain elusive.

In this study, we investigated the function of SET-24. The SET domain of SET-24 shares similarity with the catalytically inactive SET domain of MLL5 in *Homo sapiens* (HsMLL5), which belongs to the *Saccharomyces cerevisiae* (ScSET3) subfamily. This suggests the SET domain of SET-24 is catalytically inactive. Animals with *set-24* mutations, with multiple deletions in its coding sequence exhibit a Mrt phenotype at 25°C, indicating that SET-24 is required to maintain germline quiescence across generations. GFP RNAi inheritance assay demonstrated that SET-24 plays a role in the maintenance of transgenerational gene silencing. Furthermore, we found that SET-24 is a germline-specific factor localised in the nucleus. We identified HCF-1 as an interacting partner of SET-24, which is also involved in regulating TEI. Through chromatin and small RNA profiling, we found that SET-24 is required to regulate H3K4me3 and 22G-RNA levels in a subset of genes targeted by small RNA pathways. In summary, our findings reveal that SET-24, a factor with a likely catalytically inactive SET domain interacts with HCF-1 and is required at the interface between chromatin modifications and small RNA production, maintaining TEI.

## Results

### SET-24 is a member of the ScSet3 SET subfamily

The *C. elegans* genome encodes dozens of SET domain-containing proteins^20,38,39^. By aligning the SET domain sequences of these proteins and constructing a phylogenetic tree, we found that the SET domain of SET-24 is most similar to those of SET-9 and SET-26, paralogs with over 90% sequence similarity^40^ (Fig. 1a). The SET domains of SET-9 and SET-26 are likely homologous to the SET domains of SET3 in *Saccharomyces cerevisiae* (ScSET), UpSET in *Drosophila melanogaster* (DmUpSET), and MLL5 in *Homo sapiens* (HsMLL5), all of which belong to the ScSet3 subfamily of SET domains^41–43^. We also aligned the SET domain sequence of SET-24 with those of human proteins containing SET domains. Among all these proteins, the SET domain of SET-24 shares the highest similarity with those of HsMLL5 and HsSETD5 (Fig. 1b).

**Figure 1.**
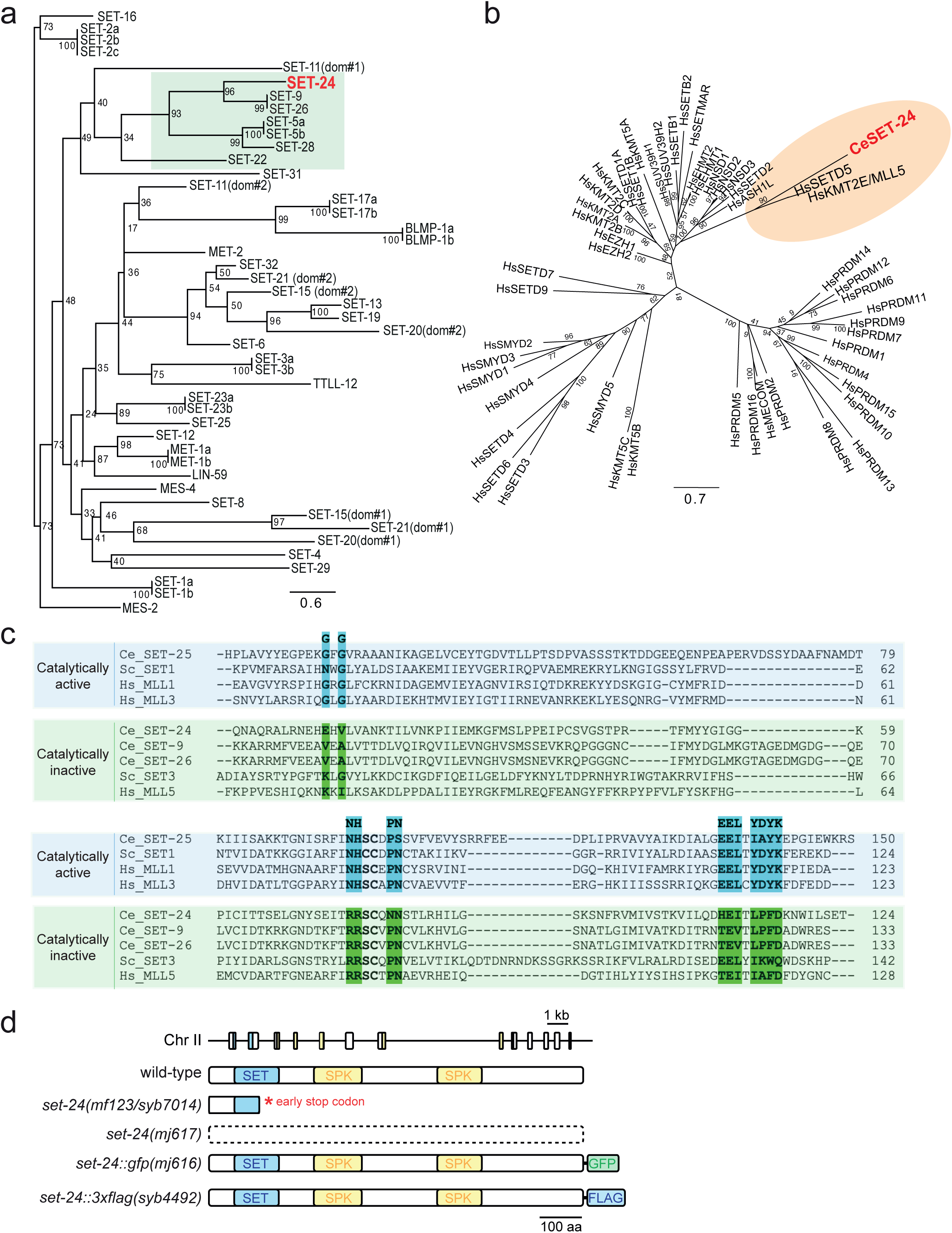
The SET domain of SET-24 is a member of the ScSet3 SET subfamily and is likely catalytically inactive. **a** Maximum likelihood phylogenetic tree comparing the protein sequences of SET domains of selected SET domain-containing genes in *C. elegans*. Values next to the tree nodes are branch supports, calculated with 1,000 ultrafast bootstrap replicates. **b** Phylogenetic tree of all human SET domains and the SET domain of *C. elegans* SET-24. Values next to the tree nodes are branch supports, calculated with 1,000 ultrafast bootstrap replicates. **c** Protein sequence alignment of the *C. elegans* SET-24 SET domain with SET domains from catalytically inactive Set3 SET subfamily and catalytically active CeSET-25, ScSET1, HsMLL1, and HsMLL3. The residues important for catalytic activity are highlighted. Ce, *Caenorhabditis elegans*; Hs, *Homo sapiens*; Sc, *Saccharomyces cerevisiae*. **d** Schematic representation of the *set-24* locus, drawn to scale, and drawings of the expected protein products corresponding to the wild-type, mutant and tagged alleles used in this study.

Members of the ScSet3 subfamily are generally considered catalytically inactive in regard to methyltransferase activity^42^. To further investigate this, we aligned the sequence of SET-24’s SET domain with archetypal active and inactive SET domain-containing proteins. Like the ScSet3 subfamily members, the SET domain of SET-24 lacks several key residues that are crucial for catalytic activity (reviewed in^42,44,45^). For instance, the asparagine-histidine (NH) motif, essential for hydrogen bonding of the methyl donor SAM and the tyrosine (Y) motif, which functions by positioning target lysine in the catalytic site, are absent in SET-24 (Fig. 1c). These similarities to the ScSet3 subfamily and the absence of conserved motifs indicate that SET-24 is a member of the ScSet3 subfamily. Besides SET-9/24/26, three other SET domains, of SET-5/22/28, lack catalytic residues and are therefore also members of the ScSet3 subfamily (Fig. S1). This suggests that the SET domains in the branch highlighted in Fig. 1a are likely catalytically inactive.

Most members of the ScSet3 subfamily possess a conserved PHD domain and a SET domain within their sequences^46^ (Fig. S2a). While the SET-24 sequence lacks the PHD domain, it contains two SPK domains (associated with SET, PHD, and Protein Kinase) of unknown function^47^ (Figs. 1d and S2a). Proteins containing SPK domains are predominantly found in nematodes, with some present in *C. elegans* (Fig. S2b). We used AlphaFold3 to predict the structures of the SPK domains (SPK1 and SPK2) in SET-24 and conducted a search for similar structures with Foldseek. The folds of the SET-24 SPK domains are very similar to the MYB-like domains of the telomere-binding proteins *C. elegans* TEBP-1 and TEBP-2, which can bind directly to double-stranded telomeric DNA sequences using their third MYB-like domain^48,49^ (Fig. S2c).

### *set-24* mutants display germline abnormalities and occasional escape from sterility

To explore the role of SET-24 in *C. elegans*, we first characterised the phenotypic impact of *set-24* mutations. To address this, we created two alleles in a wild-type N2 background: *set-24(mj617)*, an allele with a deletion of the entire coding sequence of *set-24* and *set-24(syb7014)*, a recreation of the *set-24* allele *mf123* from a previously described wild isolate that encodes a truncated version of the protein due to an early STOP codon after the 188^th^ amino acid ^35^ (Fig. 1d). When grown at 25°C, the reference N2 strain could survive and usually be maintained for an indefinite number of generations, while the nuclear RNAi-defective mutant *hrde-1* worms became sterile after 1 to 8 generations, showing a Mrt phenotype^25^. Similarly, When *set-24(syb7014)* and *set-24(mj617)* mutants were cultured at 25°C, the worms exhibited the Mrt phenotype and became sterile after 1 to 8 generations (Figs. 2a and 2b). Remarkably, certain *set-24* lines escaped the Mrt phenotype and remained fertile at 25°C for more than 20 generations. We referred to these lineages as “escapees” (Figs. 2a and 2b).

**Figure 2.**
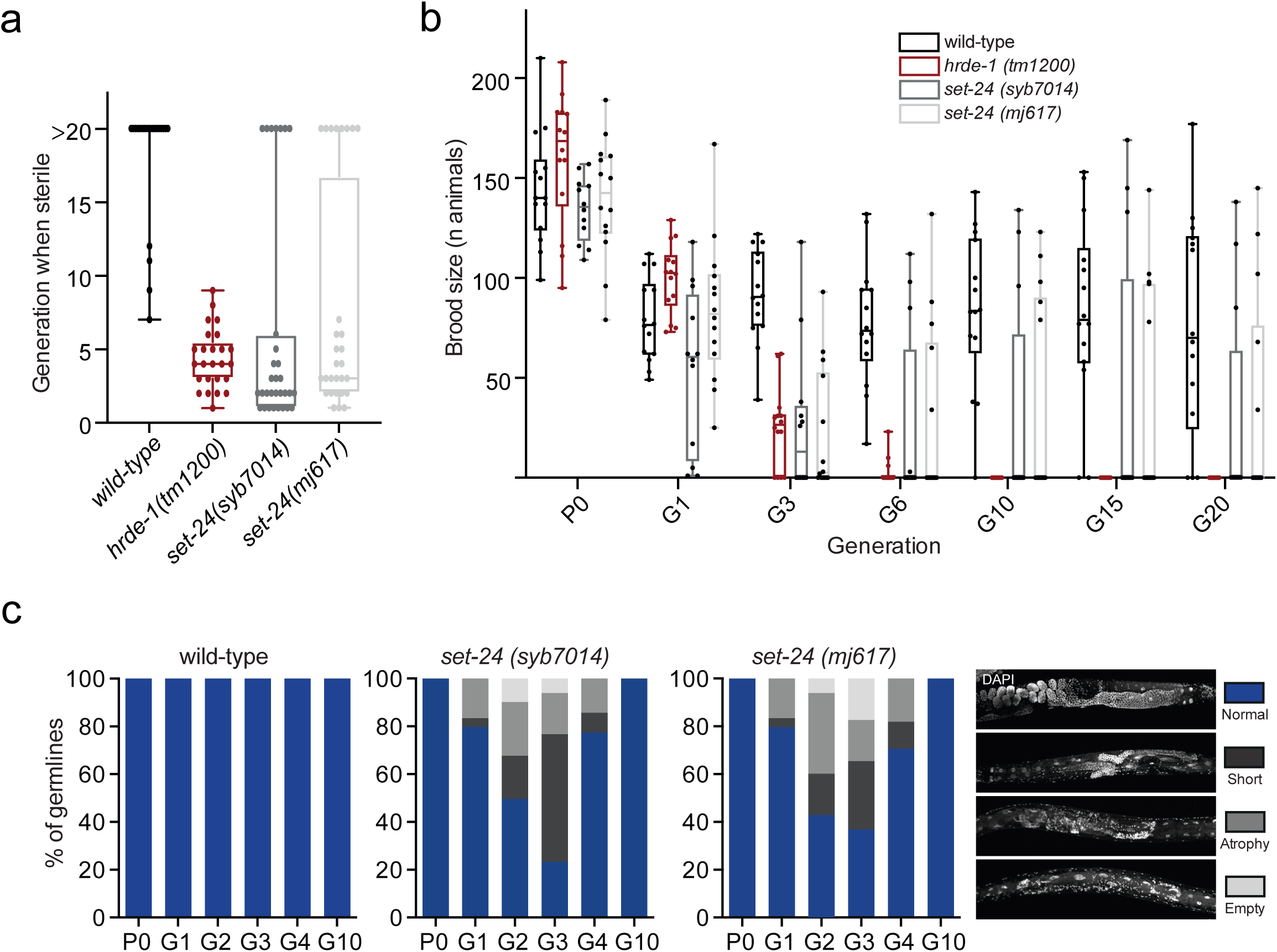
SET-24 is required for germline integrity and fertility. **a** and **b** *set-24* mutants are germline mortal (Mrt) at 25°C. **a** Boxplots showing the number of generations elapsed until animals become sterile (n>20) and, **b** quantification of the number of progeny over generations (n=15 per genotype). **c** *set-24* mutants display progressive germline degeneration under heat-stress. Quantification of the proportions of normal, short, atrophic, and empty germlines (countings on pooled progeny of 15 parents. n>30 per genotype).

After growing the worms at 25°C for several generations (G1-G3, from a parental P0), the germline of *set-24* mutant strains displayed abnormalities, such as a short or atrophied gonad, or even an empty germline, whereas the germline of wild-type worms remained normal. The percentage of *set-24* mutant worms with abnormal germline increased from the parental P0 to G3. However, beyond G4, most *set-24* mutant worms had a normal germline (Figs. 2c and S3), likely because the “escapees” became the dominant population and remained fertile in later generations. In conclusion, *set-24* mutants show a Mrt phenotype that is associated with defects in germline development, but some *set-24* lines escape sterility.

### SET-24 is required to sustain heritable RNAi

Germline immortality is subject to genetic regulation. Previous studies have demonstrated that the absence of various factors associated with nuclear RNAi and TEI contributes to germline immortality^4,16,17,22,24–26,31,32,50–53^. Additionally, several SET domain-containing genes have been implicated in TEI^27^. Thus, we aimed to investigate if *set-24* mutants exhibit defects in RNAi inheritance. We explored SET-24’s involvement in TEI using a previously described GFP::H2B transgenic strain^10^ (Fig. 3a). In this strain, GFP is consistently expressed in germline and embryo nuclei. This transgene can be silenced by feeding worms with bacteria expressing dsRNAs targeting the *gfp* sequence. Even after removing the RNAi trigger, the silencing persists in subsequent generations^10^ (Fig. 3b).

**Figure 3.**
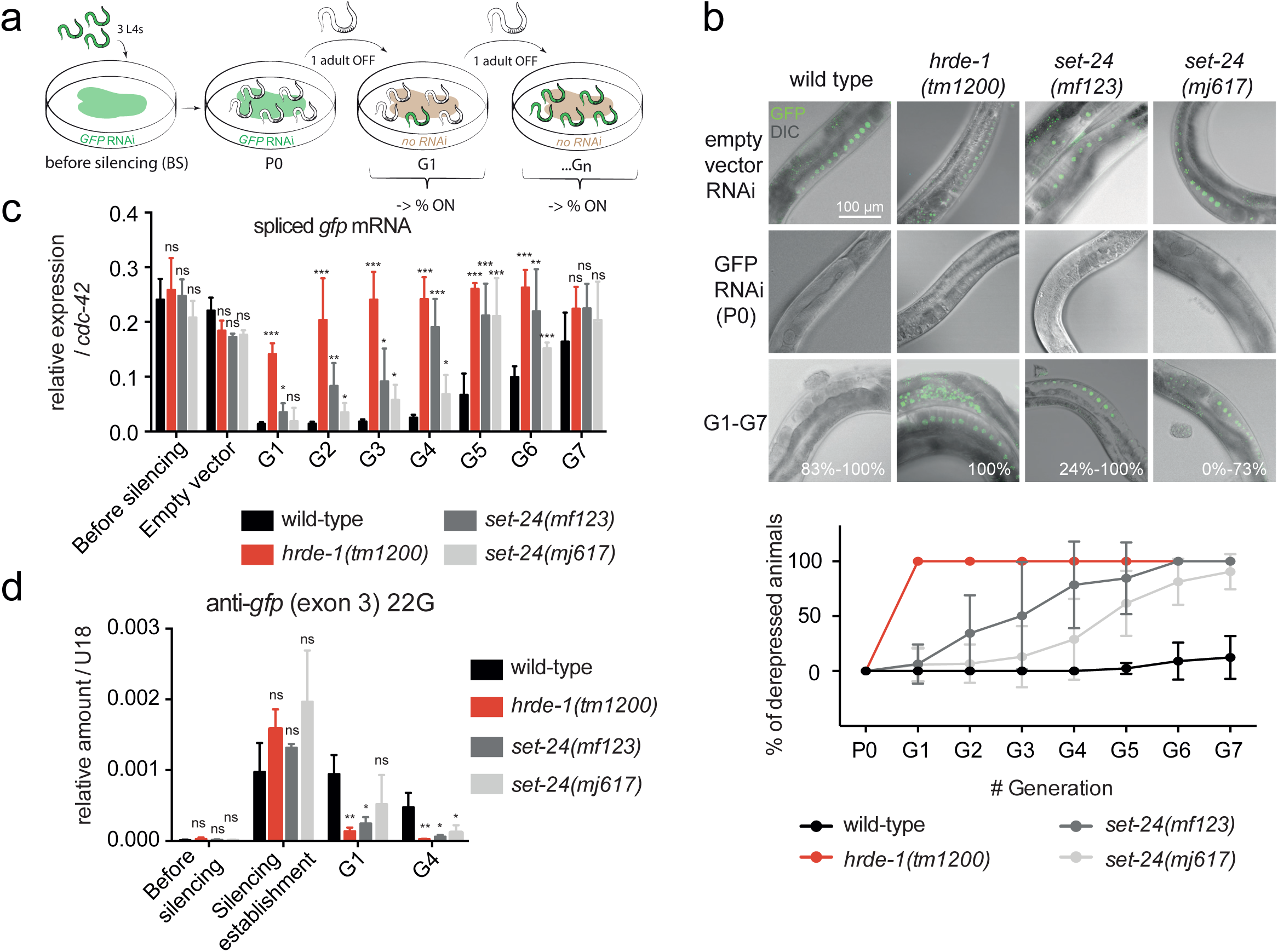
SET-24 is required for RNAi inheritance. **a** Schematic representation of the experimental procedure. **b** Top: Representative images of GFP transgene activity in *wild-type*, *hrde-1* and *set-24* mutant germlines. Scale bar = 100 μm. Bottom: Percentage of derepressed individuals in every generation. Countings on >20 animals, experiments on n=5 animal lineages per genotype, N=3 independent replicates. **c** RT-qPCR analysis of GFP transgene (spliced) mRNA expression at every generation. Means and standard deviations (SDs) over 5 animal populations are shown. St. **d** RT-qPCR measurement of the abundance of a representative anti-GFP 22G-RNA, n≥2. (c-d) Statistical significance was assessed with the unpaired t test with Welch’s correction. Stars indicate p-value, ns, not significant; * p<0.05; ** p<0.01.. Comparisons were with wild-type strain.

In worms with defective *hrde-1*, exposure to *gfp* dsRNA did not lead to heritable silencing of GFP::H2B, consistent with previous findings^25^ (Fig. 3b). In *set-24* mutant strains, the silencing gradually decreased over a few generations after the removal of the RNAi bacteria (Fig. 3b). This decline was evaluated by determining the percentage of worms expressing GFP and measuring the relative expression level of spliced GFP mRNA using quantitative polymerase chain reaction (qPCR) (Figs. 3b and 3c). Given the roles of secondary siRNAs (22G-RNAs) in heritable RNAi, we assessed the abundance of anti-GFP 22G-RNAs, which accumulate during the establishment of silencing. Anti-GFP siRNAs in *set-24* mutant worms did not persist at the same levels as in wild-type worms, one and four generations after the removal of the silencing trigger (Fig. 3d). These findings highlight the critical role of SET-24 in the maintenance of dsRNA-triggered TEI.

The piRNA pathway safeguards the germline against transposons, by recognising their transcript sand eliciting 22G-RNA biogenesis, which drive target silencing^54–56^. The piRNA sensor strain expresses an *mcherry::his-58* fusion gene with a recognition site in the 3’UTR for an abundant piRNA^57^. In a wild-type background, the sensor is silenced, but depletion of factors involved in the piRNA pathway, such as *prg-1* and *hrde-1*, leads to its derepression. To investigate the potential role of *set-24* in the piRNA pathway, we crossed *set-24* mutants with the piRNA sensor strain. In contrast to the *hrde-1* mutant, the depletion of *set-24* did not derepress the piRNA sensor (Figs. S4a and S4b). These findings suggest that *set-24* is dispensable for the initiation of piRNA-dependent silencing.

Overall, these results demonstrate that in the absence of SET-24, siRNA signals do not persist across generations and RNAi inheritance is impaired. Also, piRNA-dependent silencing is not affected by SET-24.

### SET-24 is a germline-specific factor and is localised to germline nuclei

Most TEI factors, such as the nuclear and perinuclear Argonautes, HRDE-1 and WAGO-4, respectively, are expressed in the germline^17,25,31,32^. This prompted the investigation of the expression pattern of SET-24. *set-24* mRNAs are predominantly detectable in embryo, L4, and adult stages, mirroring the mRNA expression pattern of the germline factor *pie-1* (Figs. 4a and S5a). *set-24* mRNAs were not detected in *glp-4(bn-2)* worms grown at 25°C, which lack a germline at this restrictive temperature (Fig. 4b). However, *set-24* mRNAs were still expressed in *fem-1(hc17)* and *fog-2(q71)* at the same temperature, despite the absence of sperm or oocytes respectively (Fig. S5b). These mRNA expression patterns indicate that *set-24* mRNA is confined to the *C. elegans* germline, both in spermatogenic and oogenic gonads.

**Figure 4.**
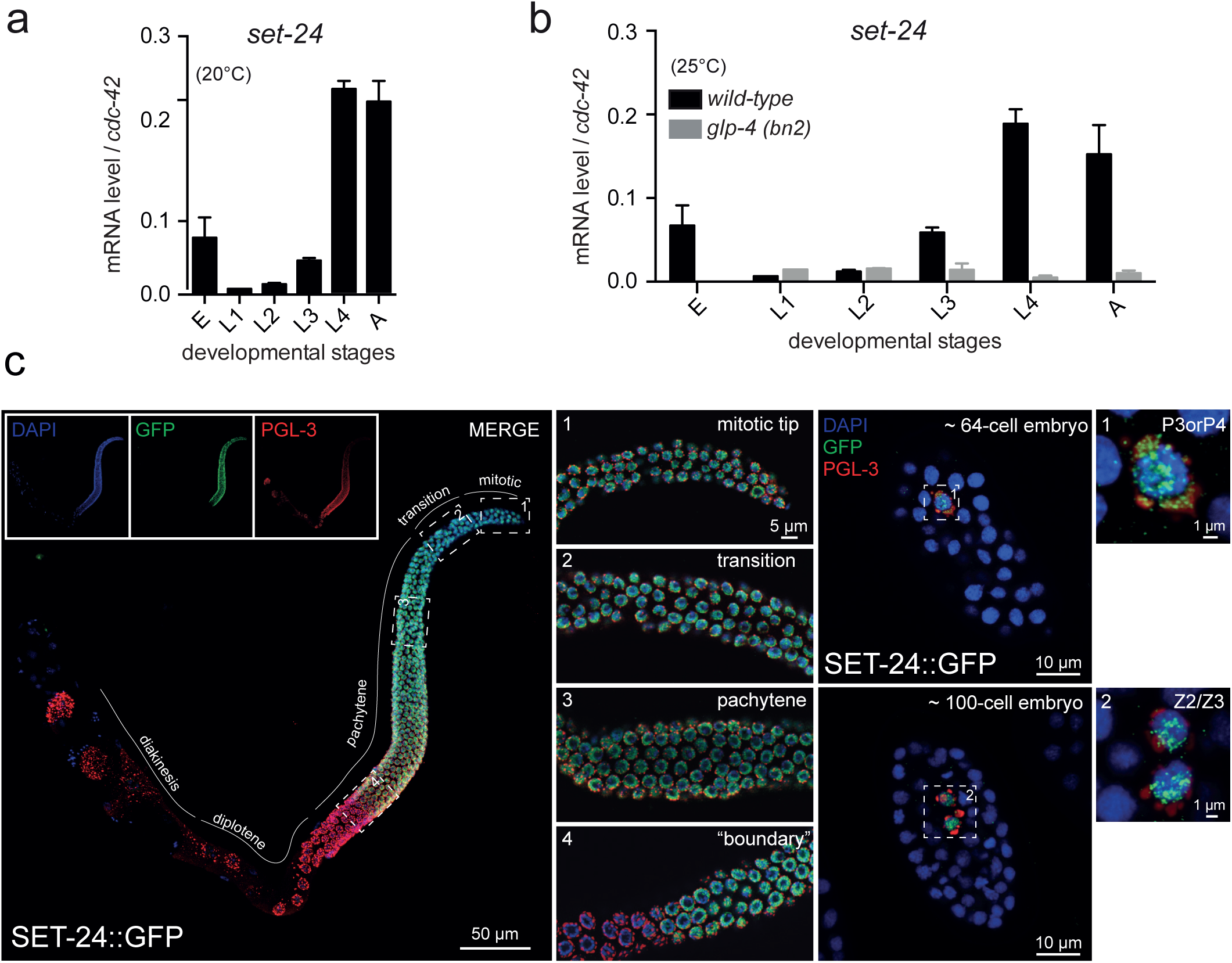
SET-24 is a germline-specific factor and is localised to germline nuclei. **a** and **b** *set-24* is germline-specific. Developmental time-course analysis of *set-24* expression by RT-qPCR in wild-type animals grown at 20°C (**a**) and in wild-type and germline-less *glp-4 (bn2ts)* animals grown at 25°C (**b**). Means and standard deviations over 3 independent experiments are shown. **c** SET-24 localises to germline nuclei. Representative immunofluorescence images of SET-24 localization (anti-GFP antibody, green) in dissected germlines of a SET-24::GFP strain. P-granules are stained with an anti-PGL-3 antibody, red, and DNA is stained with DAPI, blue. Numbered insets, zoom in on particular regions of the germline and on embryonic germ cells.

Subsequently, we examined the localization of the SET-24 protein in the germline. We created a GFP-tagged SET-24 allele (Fig. 1d). Similar to the wild-type strain, SET-24::GFP animals can survive and be maintained for numerous generations at 25°C, suggesting that the GFP at the C-terminus of SET-24 does not alter its function (Figs. S5c and S5d). SET-24::GFP can be observed in germline nuclei (Fig. 4c). Notably, unlike the germline perinuclear factor PGL-3, expressed throughout the entire gonad from the mitotic region to the diakinesis region in oocytes (Fig. S5e), SET-24::GFP was only expressed between the mitotic and pachytene regions, with a clear fade in signal at the boundary of the pachytene and diplotene regions (Fig. 4c). In embryos, SET-24::GFP was solely expressed in the nuclei of germline-lineage cells (Fig. 4c). In summary, we conclude that SET-24 is a nuclear germline-specific factor, expressed up to the pachytene region.

### The SET-24 interactor, HCF-1, inhibits the maintenance of TEI

Given the nuclear localization of SET-24, we inquired if SET-24 is able to bind to chromatin and interact with other chromatin factors *in vivo*. To investigate this, we generated a version of SET-24 endogenously tagged with 3xFLAG tag (Fig. 1d) and performed a ChIP assay using both this allele and the SET-24::GFP strain. These immunoprecipitations yielded no detectable DNA enrichment (Fig. S6a). This could be attributed to either lack of SET-24 binding to chromatin, or technical issues in our ChIP procedures.

Subsequently, we identified SET-24 binding partners through IP-MS (Immunoprecipitation and Mass Spectrometry) and Y2H (Yeast-two Hybrid) assays using YA worms expressing SET-24::3xFLAG. Host Cell Factor 1 (HCF-1), the ortholog of human HCFC1 and HCFC-2, was identified by the two assays (Figs. 5a and S6b, Supplementary tables 1 and 2). HCF-1 is a highly conserved chromatin adapter protein that recruits histone-modifying complexes to chromatin^58,59^. It has been reported that HCF-1 plays a role in longevity and stress resistance in *C. elegans*^60,61^. In both adult worms and embryos, HCF-1 is a nuclear factor expressed in both germline and soma, while SET-24 is restricted to the germline^60^ (Figs. 4c, S6c, and S6d). In the gonad, HCF-1::GFP does not show a decrease in signal at the boundary between the pachytene and diplotene regions, unlike SET-24::GFP (Fig. S6e). In the germline, no significant changes in the expression and localization of SET-24 were observed when HCF-1 is depleted, and vice versa (Fig. S6f). We propose that SET-24 and HCF-1 interact with each other in germline nuclei, but do not affect each other’s expression or localization.

**Figure 5.**
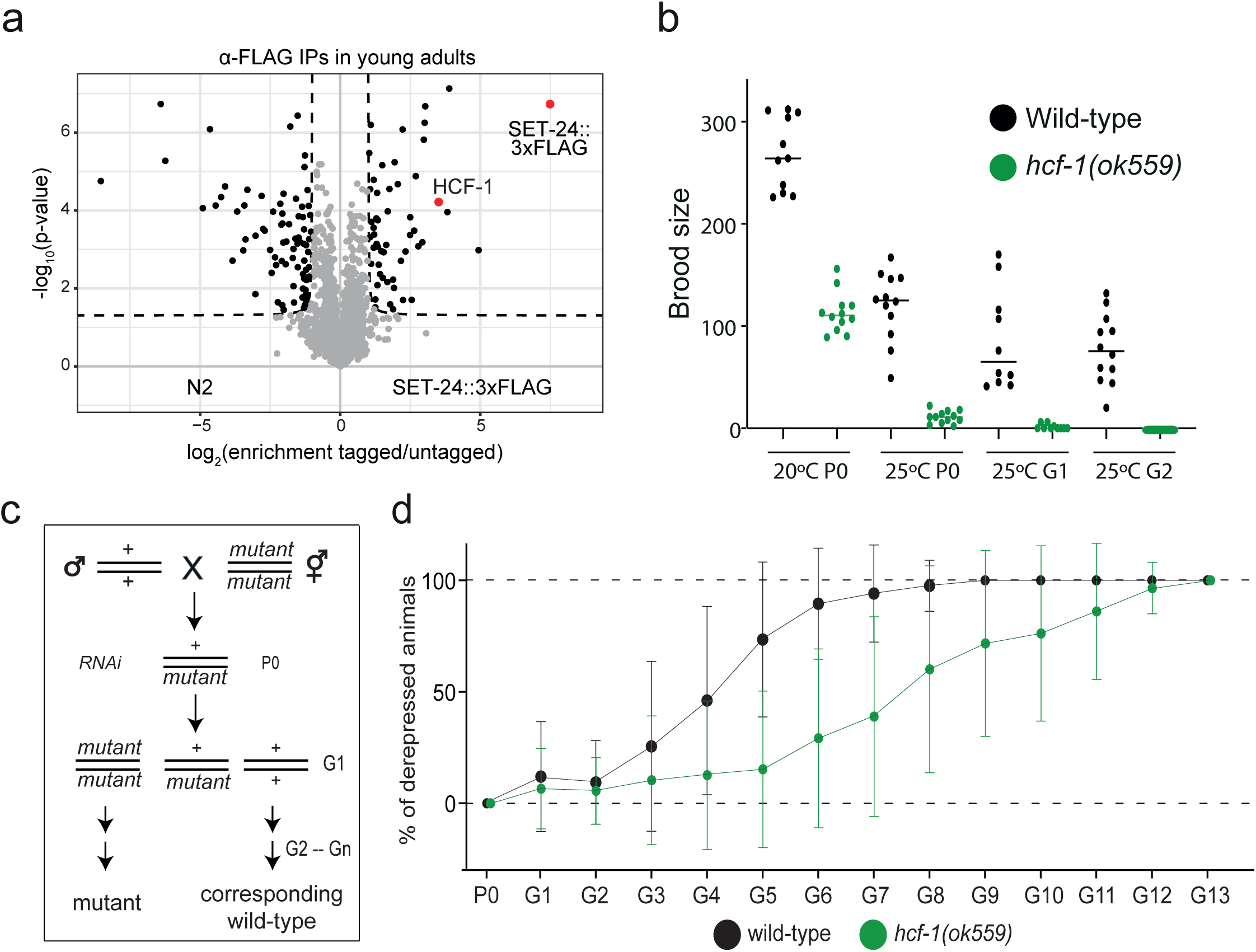
The SET-24 interacts with HCF-1, which is required for heritable RNAi. **a** Identification of SET-24 interactors via IP-MS. **b** Brood size of wild-type and *hcf-1* mutant worms at 20°C and 25°C over generations. n ≥ 10. **c** The scheme of RNAi inheritance assay for the right plot: these heterozygous individuals are fed GFP RNAi bacteria. Silenced mutants and their corresponding wildtypes, selected by genotyping PCR, are allowed to self-fertilize to produce the F1 and later generations, and are fed on HB101-seeded plates. **d** Quantification of the percentage of derepressed individuals at every generation. n > 20.

*hcf-1* mutants have approximately half the brood size of wild-type worms at 20°C and are almost sterile at 25°C (Fig. 5b), which differs from the phenotype of *set-24* mutants at 25°C (Figs. 2a and 2b). To assess if HCF-1 plays a role in TEI, we conducted a RNAi inheritance assay. Heterozygous *set-24* or *hcf-1* worms with a GFP::H2B transgene were fed *gfp* dsRNA, and the inheritance of silencing was monitored in subsequent generations. (Fig. 5c). In this experimental setup, *set-24* mutants lose silencing faster than wild-type animals (Figs. S6g and S6h), which is consistent with previous results (Fig. 3b). In contrast, it takes more generations for *hcf-1* mutants to lose silencing compared to wild-type worms (Figs. 5d and S6h). These results indicate that HCF-1 and SET-24 both mediate TEI, but display opposite phenotypes.

### SET-24 is required to modulate H3K4me3 and small RNA levels

SET domain proteins are known to influence histone modifications^62^. Human HCF-1 interacts with mixed-lineage leukemia (MLL) and the H3K4 methyltransferase Set1^59,63,64^. In *C. elegans*, HCF-1 associates with the COMPASS complex, where the H3K4 methyltransferase SET-2 serves as an essential component^65,66^. To investigate if the depletion of *set-24* affects the global levels of histone H3 methylation, we conducted a western blot assay using antibodies specific to various histone methylation marks in young adult (YA) worms. This assay was performed on worms either grown at 20°C or for three generations at 25°C. There were no observable changes in *set-24* mutants compared to wild-type worms (Fig. S7a).

Since western blotting lacks the sensitivity to detect local changes in histone modification levels, we performed a ChIP-seq assay using an H3K4me3 antibody on both wild-type and *set-24* mutant YA worms grown at 20°C. H3K4me3 is enriched at the transcription start sites (TSS) of protein-coding genes in both wild-type and *set-24* mutant worms (Fig. 6a), consistent with known patterns of this histone modification^67^. Interestingly, we observed a subtle increase in H3K4me3 signal in *set-24* mutants (Figs. 6a and 6b). This increase is restricted to genes with TSSs already marked with H3K4me3, indicating that SET-24-dependent regulation is not genome-wide and does not affect genes already decorated with H3K4me3 (Fig. 6b). We identified a subset of genes with significantly elevated H3K4me3 levels in *set-24* mutants, which we refer to as H3K4me3-enriched genes (Figs. 6a–c). Notably, this increase in H3K4me3 occurs both upstream and downstream of the TSS (Fig. S7b).

**Figure 6.**
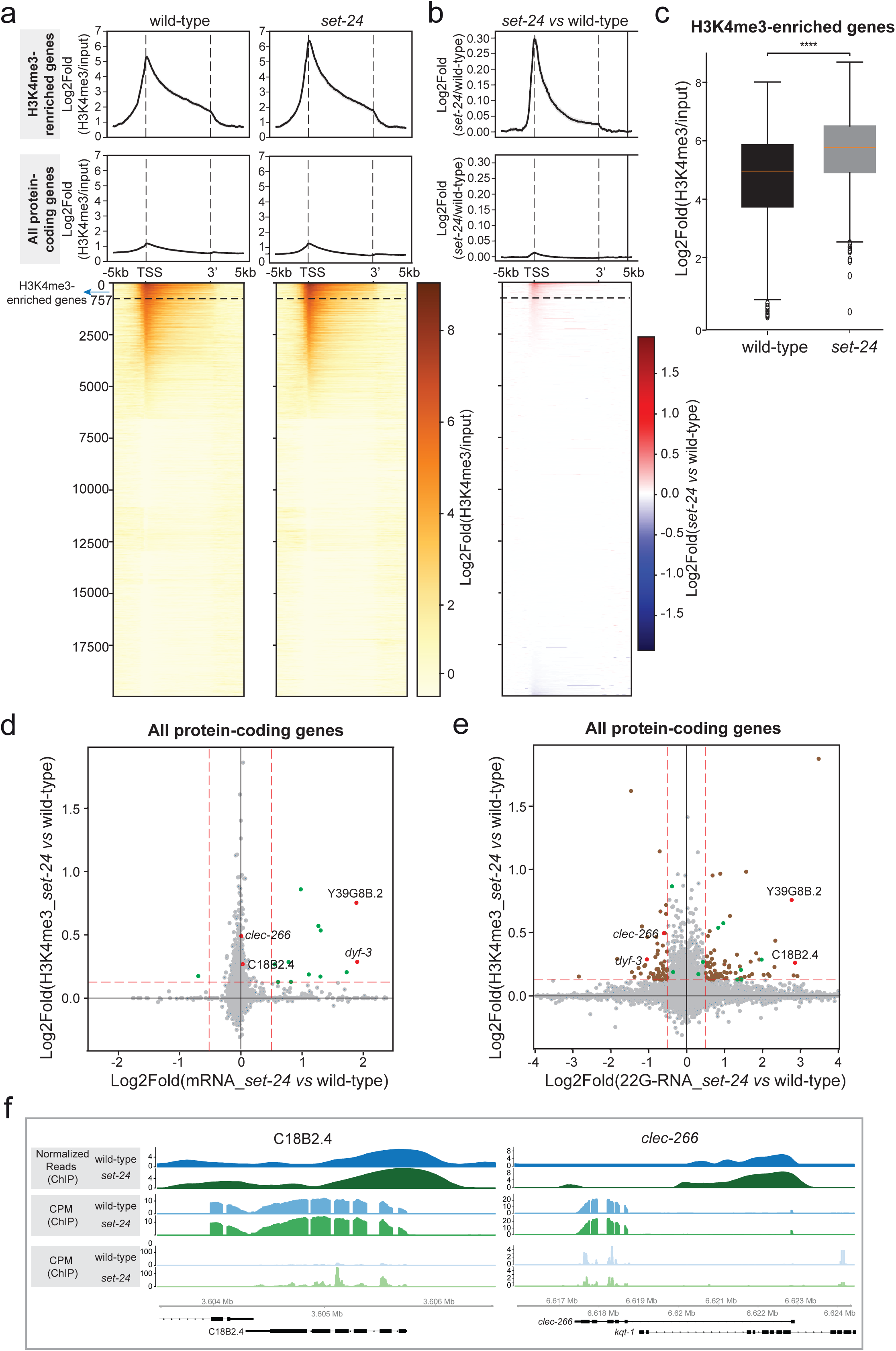
SET-24 modulates H3K4me3 accumulation and small RNA biogenesis. **a** and **b** Metagene plots and heatmaps showing H3K4me3 enrichment normalized to input for wild-type samples and *set-24* mutants across H3K4me3-enriched genes and all protein-coding genes (**a**) and the log₂-normalized fold-change between *set-24* mutants and wild-type samples (**b**). A subset of H3K4me3-enriched genes that contribute to 80% of the total increased enrichment in *set-24* mutants is labelled in the heatmaps. **c** Comparison of the average H3K4me3 enrichment around the TSS (±500 bp) for wild-type samples and *set-24* mutants across H3K4me3 enriched genes. Statistical significance was assessed using a paired *t*-test. Asterisks indicate: ****p < 0.001. **d** and **e** Scatter plots correlating the increased H3K4me3 enrichment around the TSS and mRNA levels of protein-coding genes (**d**) or 22G RNA levels across all protein-coding genes(**e**). These genes were selected from those with H3K4me3 increase that also exhibited a log₂-normalized increase of ≥0.5 in mRNA levels or a log₂-normalized change of ≥0.5 in 22G RNA levels. **f** H3K4me3 enrichment, mRNA and 22G-RNA levels of the selected genes (C18B2.4 and *clec-266*) in wild-type and *set-24* mutant strains.

H3K4me3 is generally considered a histone mark associated with transcriptional activation^68^. To assess if the observed increase in H3K4me3 levels influences gene expression, we performed mRNA sequencing on wild-type and *set-24* mutant YA worms and analysed the transcriptional changes across all genes (Supplementary table 3). We identified 12 genes with both H3K4me3 enrichment and mRNA upregulation in *set-24* mutants (Fig. 6d and Supplementary table 4). These genes are also regulated by components of the COMPASS complex, SET-2 and WDR-5, as determined through a WormExp enrichment analysis^69^ (Supplementary table 5), further suggesting a role for SET-24 in H3K4me3 regulation through HCF-1 and its interaction with COMPASS. However, the majority of H3K4me3-enriched genes did not exhibit increased transcription in *set-24* mutants (Fig. 6d).

Given the role of SET-24 in the maintenance of heritable RNAi and inheritance of 22G-RNAs (Fig. 3b-d), we sought to investigate the potential link between SET-24 and 22G-RNA regulation. To do so, we sequenced small RNAs in wild-type and *set-24* mutants and analysed 22G-RNAs mapping to protein-coding genes. We identified genes with significant changes in 22G-RNA expression in *set-24* mutants relative to wild-type (Supplementary table 6), and classified them as *set-24*-regulated 22G-RNA targets. 22G-RNAs can be subdivided into distinct subpopulations, according to the Argonaute protein they associate with, the factors required for their biogenesis, and the genes targeted^70^. We found that the *set-24-*regulated 22G-RNA targets overlap with WAGO- and Mutator-regulated genes (Fig. S7c), suggesting SET-24 is required for silencing of a subset of endogenous WAGO and Mutator targets.

Next, we examined the correlation between H3K4me3 enrichment and *set-24*-regulated 22G-RNA targets (Fig. 6e). In total, 131 out of 757 H3K4me3-enriched genes in *set-24* mutants exhibited disrupted 22G-RNA levels (73 upregulated and 58 downregulated), whereas only 13 out of 757 showed altered transcriptional levels (including 12 upregulated and 1 downregulated, Figs. 6d and 6e, Supplementary table 4). Among the H3K4me3-enriched genes, few (e.g.*Y39G8B.2* and *dyf-3*) exhibited both increased transcription and disrupted 22G-RNA expression (Fig. S7d). However, a larger subset, including *C18B2.4* and *clec-24*, showed disrupted 22G-RNA levels without corresponding transcriptional changes (Figs. 6e and 6f). These results suggest that SET-24 is dedicated to maintaining 22G-RNA levels.

Taken together, these results suggest that SET-24 modulates H3K4me3 and 22G-RNA levels in a subset of WAGO/mutator target genes.

## Discussion

In this work, we provide insights on the roles of SET-24, a likely catalytically inactive SET domain-containing factor specifically expressed in germline nuclei. SET-24 maintains germline integrity across generations and is, together with its interactor HCF-1, required for heritable RNAi. We will discuss below how the imbalance of H3K4me3 and 22G-RNA levels observed in *set-24* mutant animals may be relevant for the maintenance of epigenetic memory.

The SET domain superfamily consists of multiple subfamilies, one of which is the ScSET3 family. ScSET3 family members are characterized by similarities in domain structure, sequence, and biological function to the Set3 protein of *S. cerevisiae*^71^. Although these members are unlikely to possess catalytic methyltransferase activity, they play essential roles in various biological processes and are associated with epigenetic regulation^46,72^. However, mechanistic insight into the function of ScSET3 subfamily members is lacking. Notably, most ScSET3 family proteins, except for SETD5 in zebrafish, mouse, rat, and human, contain a PHD domain—a chromatin reader that recognizes methylated H3 lysine 4 (H3K4)^46,73,74^. Despite its role in the maintenance of H3K4me3 levels, SET-24 lacks a PHD domain and is therefore unlikely to recognise H3K4 directly. Some members of the ScSET3 subfamily regulate gene expression by recruiting histone deacetylases (HDACs) to specific genomic regions^46,61,71,75^. However, our IP-MS and yeast two-hybrid assays did not identify any HDACs as interacting partners of SET-24 (Supplementary tables 1 and 2), arguing against a direct interaction with HDAC complexes.

Unlike other ScSET3 subfamily members that possess PHD domains, SET-24 contains two SPK domains. The biological and biochemical functions of these SPK domains remain unclear. Interestingly, the SPK domains of SET-24 share structural similarity with MYB-like domains found in TEBP-1 and TEBP-2 (Fig. S2c). One of three MYB-like domains in TEBP-1 and TEBP-2 directly binds to double-stranded telomeric DNA sequences^49^. Although we were unable to immunoprecipitate SET-24 together with chromatin, exploring the DNA-binding potential of the SET-24 SPK domains using alternative approaches could provide valuable insights. SPK domain-containing proteins are predominantly found in nematodes, with several identified in *C. elegans* (Fig. S2b). We aligned SPK domains in *C. elegans* and found that the SPK2 domain in SET-24 is closely related to the SPK domain in OGR-2, which is known to influence progression of oogenesis in *C. elegans*^76^ (Fig. S2b). This similarity suggests that SET-24 might regulate germline development through its SPK domains. Interestingly, another SET domain-containing protein, SET-5, also contains an SPK domain, and its SET domain is very similar to that of SET-24 and is likely catalytically inactive (Figs. 1a, S1, and S2b). It will be important to determine if SET-24 and SET-5 have overlapping and/or redundant functions. The relatively small number of genes with associated H3K4me3 and 22G-RNA imbalance, may be explained by redundancy with other SET domain-containing proteins.

Additional SET domain-containing proteins highly likely to have redundant functions with SET-24 are SET-9 and SET-26. These three SET domain-containing proteins are required for heritable RNAi, germline immortality at 25°C, and interact with HCF-1^61,77^. While we identified HCF-1 as a SET-24 interactor (Figs. 5A and S6b), IP-MS and Y2H analyses of the interactome of SET-24 did not detect SET-9 or SET-26. Likewise, SET-24 was not identified in the interactome of SET-9/26, suggesting that SET-24 does not interact directly with these other SET domain-containing proteins. Of note, SET-9 and SET-26 bind to HCF-1 through a conserved motif^61,78^ (Fig. S2A), which is absent in SET-24, implying that SET-24 may interact with HCF-1 in a different manner.

The subcellular localisation and expression pattern of these SET domain-containing proteins are also consistent with functional redundancy. Much like HCF-1, SET-9/26 and SET-24 are all expressed in germline nuclei^41,60^ (Figs. 4c and S6c-e), although HCF-1 and SET-26 are present in somatic nuclei as well. The expression pattern of SET-26 and HCF-1 is consistent with additional regulatory roles outside the germline. Indeed, SET-26 was shown to modulate the lifespan of *C. elegans* in conjunction with HCF-1 by binding to H3K4me3 in somatic cells^61^. In the germline, SET-24 displays a pattern of expression from the mitotic tip to the pachytene-diplotene boundary region (Fig. 4c), indicating that SET-24 might have a regulatory role up until the pachytene-diplotene transition and affect normal meiotic progression. In fact, this is a germline region where chromatin changes required for meiotic progression, chromosomal organization and recombination have been previously described^79–81^. Conversely, HCF-1 and SET-9/26 are expressed throughout the gonad^41,60^ (Fig. S6d). Therefore, in mitotic and early meiotic stages, SET-24 is co-expressed with SET-9/26 in germline nuclei, where these factors associate with HCF-1 and may have partial redundant regulatory roles. At this point, we cannot exclude the alternative that SET-24 competes with SET-9/26 for HCF-1 binding.

Small RNA-driven TEI consists of three steps: initiation, establishment, and maintenance^18^. Our RNAi inheritance assays show that SET-24 and HCF-1 act exclusively in the maintenance step. This is unusual, as most TEI factors, such as HRDE-1, WAGO-4, and ZNFX-1, play roles in both the establishment and maintenance phases, whereas SET-25 and SET-32 are primarily involved in the establishment^27^. SET-24 and MET-2 are the only SET domain-containing proteins with an exclusive role in the maintenance stage^22^. Notably, SET-24 and HCF-1 contribute distinctly to the maintenance of heritable RNAi, as *set-24* and *hcf-1* mutants display opposite phenotypes: shortened and extended transgenerational RNAi-driven silencing, respectively (Figs. 3b, 5d, S6g and S6h). This is likely explained by the myriad roles played by HCF-1 in the context of different protein complexes, compared to more specialised SET-24. Besides interaction with SET-24, HCF-1 interacts with the H3K4 methyltransferase SET-2 and the COMPASS complex, other chromatin remodelling complexes, and with the H3K4me3 readers SET-9 and SET-26^41,59–61,63–66,82,83^. These interactors have distinct effects on TEI and may explain the role of HCF-1 in inhibiting TEI. For example, while SET-9/26 influence TEI^77^, SET-2 is not required for the process^84^. This role of HCF-1 in extending RNAi inheritance is not unprecedented, as similar roles have been reported for HERI-1, MET-2, and LOTR-1^19,22,85^. Whether HCF-1 is acting in concert with HERI-1, MET-2, or LOTR-1 remains to be determined. Further research is needed to better understand how SET-24 and HCF-1 regulate TEI.

Our data suggests an imbalance of H3K4me3 and 22G-RNA levels caused by *set-24* mutation (Figs. 3d and 6). We propose that this imbalance, together with our phenotypic data, reflects a role of SET-24 in the maintenance of epigenetic memory across generations. We found that H3K4me3 levels of the TSSs of 757 genes are regulated by SET-24, with an upregulation observed in *set-24* mutants that is not consistently accompanied by transcriptional activation of these genes. However, a larger subset of 131 genes with SET-24-dependent H3K4me3 regulation tend to have deregulated 22G-RNA levels. Genes with deregulated 22G-RNA levels are targets of silencing 22G-RNA pathways in wild-type (Fig. S7c). The relatively small number of genes affected may be due to possible redundancy with other germline-expressed SET domain-containing proteins, as discussed above. We investigated the levels of H3K4me3 given the association of SET-24 with HCF-1, which is a cofactor of H3K4me3-directing COMPASS complex. However, the lack of transcriptional upregulation of SET-24-dependent H3K4me3-enriched genes may be due to other repressive chromatin modifications that were not profiled in our study, such as H3K9me3. As H3K9me3 is a mark associated with the WAGO and mutator 22G-RNA silencing pathways targeting SET-24-dependent genes (Fig. S7c), we postulate that the interplay between H3K4me3 and H3K9me3 is disrupted in *set-24* mutants and affects the maintenance of the chromatin environment adequate for the maintenance of gene silencing across generations. Lack of sustained 22G-RNA biogenesis may in turn contribute to further destabilisation of the chromatin environment at these loci. Further genetic and biochemical dissection of these processes is required to understand how SET-24 and other SET domain-containing proteins define the chromatin landscape and affect 22G-RNA biogenesis in the germline.

The *set-24* allele was originally identified in wild *C. elegans* isolates with a mortal germline^35^. What possible advantage, if any, could such a mutant allele confer in the wild? Perhaps the answer lies in the “escapees” identified in the fertility assays across generations (Fig. 2). The Mrt phenotype is reversed in particular animal lineages, which presumably develop a compensatory response. Interestingly, escape from the Mrt phenotype also occurs occasionally in *set-9* and *set-26* single mutant lines^77^. The “escapee” phenotype could reflect a compensatory response from redundant germline-expressed SET domain-containing factors.

Reversibility of the Mrt phenotype is not unprecedented, for example upon exposure to specific stimuli, such as temperature shifts and different bacterial diets^34,35^. A provocative alternative hypothesis consists in the establishment of secondary chromatin-state and small RNA-based epimutations, some of which may be advantageous. In line with this hypothesis, prominent 22G-RNA-based epimutations in WAGO targets have been documented^86^. Therefore, natural genetic variation may disrupt epigenetic processes, like in the wild-isolated strain defective for *set-24*, leading to secondary epimutations on the chromatin state and small RNA regulation. These epimutations could help wild worm populations deal with and adapt to environmental fluctuations, supporting the essential roles played by small RNA pathways in pathogen and environmental stress responses^87–92^.

## Materials and methods

### Evolutionary and structural analysis of SET and SPK domains

As the hidden Markov model (HMM) for SET domains within Pfam-A.hmm (version 3.1.b2, Feb 2015)^93^ was not sensitive enough to retrieve the SET domain of SET-24 of *C. elegans*, we first constructed an alternative HMM. To do so, we first downloaded the protein sequences of the InterPro^94^ SET domain (IPR046341), and the metadata with the associated domain coordinates, filtering for all human SET domains, as well as the catalytically inactive SET proteins of *Drosophila melanogaster* (UpSET) and *Saccharomyces cerevisiae* Set3. Then, we used the domain coordinates to trim all these protein sequences, keeping only the sequences of the SET domains, which were used as input for multiple sequence alignment with MAFFT v7.475, using option --auto^29,95^. Then, the HMM profile was built with this alignment using hmmbuild of the HMMer package (v3.3, hmmer.org). The HMM profile was used to search the entire *C. elegans* proteome (Wormbase ParaSite, version WBPS16)^96^ for SET domains with hmmsearch of the HMMer package (version 3.3, hmmer.org), with option --nobias. The *C. elegans* SET domains were aligned with MAFFT v7.475^95^, using option --auto (model chosen L-INS-i). The resulting alignment was input to infer a maximum likelihood phylogenetic tree with IQ-TREE v2.1.2^97^ with 1000 bootstraps (option -B 1000)^98^. LG + G4 was the best fit model. We created an additional phylogenetic tree using the abovementioned human SET domains, plus the SET domain of *C. elegans* SET-24. Alignments were conducted with MAFFT (L-INS-i was the chosen model), and a tree was constructed with IQ-TREE (LG + G4 was the best fit model) as above.

We obtained the sequences of all proteins with an SPK domain (IPR006570), and the coordinates of the SPK domains from InterPro^94^. Sequences were trimmed according to the SPK coordinates to leave only the SPK sequence. The SPK domains were subsequently aligned with MAFFT v7.475^95^, using option --auto (model chosen L-INS-i), and a maximum likelihood tree was constructed with IQ-TREE v2.1.2^97,98^ with option -B 1000, and with LG + G4 as the best fit model.

The structure of SET-24 was predicted with AlphaFold3^99^, and the regions corresponding to the annotated SPK domain coordinates were extracted and used as input in Foldseek^100^. The MYB domains of TEBP-1 and TEBP-2 were amongst the top hits identified by Foldseek. We used AlphaFold3^99^ to predict the structures of TEBP-1 and TEBP-2, and extracted the structures corresponding to their MYB domains, according to previously defined domain annotations^49^ and predicted secondary structure elements. ChimeraX^101^ was used to visualise predicted models and perform structural alignments.

### Strains

The Bristol strain N2 was used as the standard wild-type strain. All strains were grown at 20°C unless otherwise specified. The strains used in this study are listed in Supplementary table 7.

### Construction of transgenic strains

Wild-type YA animals (N2 strain) were injected with a mixture of target gene HR repair template (IDT oligos) (1 mg/ml), target gene CRISPR crRNA (Dharmacon) (8 mg/ml) and His-Cas9 (in-house bacterial purification) (5 mg/ml) dissolved in injection buffer (10 mM KCL; 10 mM Tris–HCl at pH: 8.0). F1 animals were singled, allowed to produce homozygous F2 progeny by selfing and genotyped. All final strains were checked by Sanger sequencing and outcrossed twice. The *set-24 (syb7014) and the set-24::3xflag (syb4492)* strains were made by SunyBiotech. The *set-24 (mj617) and set-24::gfp(mj616)* strains were made by standard CRISPR/Cas9 methods^102^. Fot the *set-24::gfp (mj616)* strain, the homologous repair template containing the GFP sequence generated by PCR from the AP625 plasmid (Addgene), and a guide RNA, TGATCATTTCGATGACAACG, were used. For the *set-24 (mj617)* strain, two guide RNAs, TGATCATTTCGATGACAACG and ACAGCCGGTGAGAATGTGTT, and the repair temple, AAAAAAAAACAAAGGAGAGATGCATTAACTTGTAAGAAAATAATATTATCATTGAACATCCGGCGAATTTTGGAGAAAACTGGTGGTTTCGTTGAAATCCATGATTGTTCTTGTGAAATTATACACTGAAAATAAATATTTATATGTATTACTTATTTTTAAATATTTAGT, were used.

### Germline mortality assay

Worms were maintained at 20°C prior to the start of the experiment. At least 10 L4-stage individuals per genotype, designated as the P0 (parental) generation, were transferred to HB101-seeded plates at 25°C and allowed to produce offspring. At each generation, one L3 larva per replicate per genotype was transferred to a new plate to produce the next generation. The number of sterile animals at each generation was recorded.

### Brood size assay

L3 worms were individually placed onto fresh NGM plates. The number of progeny was counted in each generation over the course of three consecutive days.

### Transgenerational memory inheritance

One L4 larva per genotype was plated on either GFP RNAi-expressing bacteria or empty vector L4440 bacteria. G1 animals were examined under a fluorescence microscope, and one silenced animal per replicate per genotype was transferred onto plates seeded with standard HB101 bacteria. At each generation, a single silenced animal was isolated from each plate to produce the next generation, while the remaining adult progeny were analysed under a fluorescence microscope. At least 20 animals per replicate per genotype were counted at each generation unless otherwise specified. Germline nuclear GFP fluorescence was scored as “on” or “off.” Representative images were captured using a Leica SP8 confocal fluorescence microscope at 40X magnification.

### RNA extraction and real-time quantitative PCR

Total RNA was extracted using TRIzol reagent (Ambion, Life Technologies) and treated with Turbo DNase Kit (Invitrogen) according to the manufacturer’s instructions. 500 ng of total RNAs per sample were reverse-transcribed with random hexamers (Invitrogen) at 50°C for 1 hour using Superscript III (Invitrogen). Reactions lacking reverse transcriptase were systematically run in parallel as negative controls. Real-time quantitative PCR was performed on 1 ul of diluted (1/5) RT reactions using SYBR Green kit (Life Technologies) on a OneStepPlus thermocycler (Thermo Fisher). All samples were run in duplicates and expression levels normalized to the reference gene *cdc-42* according to the ΔΔCt method^103^.

### 22G-RNA real-time quantitative PCR

50 ng of total RNA were reverse-transcribed with a TaqMan Small RNA Assay Kit (Thermo Fisher) containing a gene-specific RT primer and a TaqMan MicroRNA Reverse Transcription Kit (Thermo Fisher), according to the manufacturer’s instructions. Real-time quantitative PCR was performed on 1 ul of diluted (1/5) RT reactions using custom-made TaqMan probe (GUGUCCAAGAAUGUUUCCAUCU), TaqMan Universal Master Mix No AmpErase UNG (Life Technologies) on a OneStepPlus thermocycler (Thermo Fisher). All samples were run in triplicates. Expression levels were normalised to the reference gene U18 (#001764, TaqMan) according to the ΔΔCt method^103^.

### Whole-mount DAPI staining

At every generation, synchronised adult worms were collected in M9, fixed in 70% ethanol for 1 hour, centrifuged and washed twice with 0.1% Tween-20/PBS (PBST) and strained in 100 mg/mL DAPI/PBST for 30 minutes at room temperature on a rotating wheel. Worms were then washed twice with PBST, pipetted onto a microscope slide and mounted in Vectashield. Worms were classified in categories and manually counted (at least 50 animals per genotype per generation were counted) under a fluorescence microscope. Single-plan representative images were taken on a SP8 confocal fluorescence microscope (Leica) at 63X magnification.

### Immunofluorescence staining

Worms were picked on glass slides and manually dissected to extrude the germlines. Germlines were subsequently freeze-cracked on dry ice and fixed in cold 100% methanol for 20 min. Fixed slides were then washed in 0.05% Tween/PBS and incubated with diluted (1:1,000) anti-GFP (Abcam #ab290) and diluted (1:1,000) anti-PGL-3 (a gift by Susan Strome) antibodies overnight. The following day, slides were washed in 0.05% Tween/PBS and incubated with diluted secondary antibodies for 1 h at room temperature and mounted in Vectashield with DAPI. Representative images were taken on a Leica SP8 confocal microscope with a 63X oil objective and 4X digital magnification. Single-plan images are shown.

### SET-24::3XFLAG immunoprecipitation and mass spectrometry

Procedure was conducted as previously described^19,48^. Wild-type N2 and SET-24::3xFLAG animals were grown at 20°C in HB101 high-density plates synchronised by bleaching and overnight hatching of L1s in M9 buffer. L1s were plated and grown at 20°C for 51-55 hours, until the YA stage. At this stage, worms were washed off plates, washed 3-4 times in M9 buffer, washed one last time with deionised water, and snap-frozen on dry ice. To prepare extracts, worm samples were thawed and mixed 1:1 with 2x Lysis Buffer (50 mM Tris/HCl pH 7.5, 300 mM NaCl, 3 mM MgCl2, 2 mM DTT, 0.2 % Triton X-100, and complete EDTA-free Mini protease inhibitors, Roche #11836170001). Lysis was subsequently performed by sonication in a Bioruptor Plus (Diagenode, on high level, 10 cycles of 30 seconds on and 30 seconds off). After sonication, the samples were centrifuged at 21,000 x *g* for 10 min to pellet cell debris, and the supernatant was transferred to a fresh tube. Protein concentrations were determined with Bradford Protein Assay (according to manufacturer’s instructions, Bio-Rad, #5000006). IPs were prepared in quadruplicates for each strain used. 30 µl of Dynabeads Protein G (Invitrogen, #10003D) were used per IP and washed three times with 1 ml Wash Buffer (25 mM Tris/HCl pH 7.5, 300 mM NaCl, 1.5 mM MgCl2, 1 mM DTT, and complete EDTA-free Mini protease inhibitors, Roche #11836170001). The beads were resuspended in Wash Buffer and combined with to 2 mg of complete protein extract, for a total volume of 500 µl. Finally, 2 µg of anti-FLAG antibody (Sigma-Aldrich, #F1804) were added, and the samples were incubated for 3h30m, rotating at 4 °C. After the incubation, the samples were washed five times with 1 ml Wash Buffer, followed by bead resuspension in 1x LDS/DTT, and boiling at 95 °C for 10 min.

IP samples were boiled at 70°C for 10 minutes and separated on a 4–12% gradient Bis-Tris gel (Thermo Fisher Scientific, #NP0321) in 1x MOPS (Thermo Fisher Scientific, #NP0001) at 180 V for 10 minutes. Then, samples were processed separately, first by in-gel digestion, followed by desalting with a C18 StageTip^104,105^. Afterwards, the digested peptides were separated on a heated 50-cm reverse-phase capillary (75 μm inner diameter) packed with Reprosil C18 material (Dr. Maisch GmbH). Peptides were eluted along a 90 min gradient from 6 to 40% Buffer B (see StageTip purification) with the EASY-nLC 1,200 system (Thermo Fisher Scientific). Measurement was done on an Orbitrap Exploris 480 mass spectrometer (Thermo Fisher Scientific) operated with a Top15 data-dependent MS/MS acquisition method per full scan.

All raw files were processed with MaxQuant^106^(version 1.6.5.0) and peptides were matched to the *C. elegans* Wormbase protein database (version WS269) including *E. coli* sequences (ASM1798v1). Raw data and detailed MaxQuant settings can be retrieved from the parameter files uploaded to the ProteomeXchange Consortium via the PRIDE repository, accession number PXD057349. Data analysis was completed in R.

### Western blotting

Synchronized YA stage worms either growing at 20°C or three generations at 25°C were harvested and washed three times with M9 buffer before being frozen at −80°C. Worm proteins were extracted by heating the samples at 95°C for 10 minutes in 1× protein dye (62.5 mM Tris pH 6.8, 10% glycerol, 2% SDS, 5% β-mercaptoethanol, 0.2% bromophenol blue). Samples were then spun at high speed for 1 minute to remove insoluble components, and the supernatant was quickly transferred to a new tube on ice. The samples were either immediately loaded onto a gel or stored as at –80°C.

Proteins were separated by SDS-PAGE on gradient gels (10% separation gel, 5% spacer gel) and transferred onto a Hybond-ECL membrane. After washing with 1× TBST buffer (Sangon Biotech, Shanghai) and blocking with 5% milk-TBST, the membrane was incubated overnight at 4°C with primary antibodies (listed below). The next day, the membrane was washed three times for 10 minutes each with 1× TBST, followed by incubation with secondary antibodies at room temperature for 2 hours. After three additional 10-minute washes with 1× TBST, the signal was visualized.

The primary antibodies used were β-actin (Beyotime, AF5003), H3 (Abcam, ab1791), H3K4me3 (Abcam, ab8580), H3K9me1 (Abcam, ab9045), H3K9me2 (Abcam, ab1220), H3K9me3 (Millipore, 07-523), H3K23me2 (Active Motif, 39653), H3K23me3 (Active Motif, 61499), H3K27me3 (Millipore, 07-449), and H3K36me3 (Abcam, ab9050). The secondary antibodies used were goat anti-mouse (Beyotime, A0216) and goat anti-rabbit (Abcam, ab205718).

### ChIP and ChIP-seq analysis

ChIP protocol was based on a previously published protocol^107^. Synchronized worms at the YA stage were collected in M9 buffer and rapidly frozen in liquid nitrogen to create worm pellets. The worm pellets were transferred to a metallic grinder that had been pre-cooled in liquid nitrogen for 5 minutes. The worms were ground until broken into small pieces while keeping the nuclei intact. The resulting powder was transferred into a cold 50 mL Falcon tube. The crosslinking was performed on a 40ml PBS solution containing 1% formaldehyde by shaking at room temperature for 8 minutes. Then, 4.6 mL of 1.25 M glycine was added to quench the reaction, and the mixture was gently shaken for another 8 minutes at room temperature. The sample was washed twice in PBS with protease inhibitor (PI, cOmplete Tablets, Mini EDTA-free, *EASYpack*, REF #04693159001) and once in FA buffer (50 mM Hepes/KOH pH 7.5, 1 mM EDTA, 1% Triton X-100, 0.1% sodium deoxycholate, and 150 mM NaCl) with PI. The worms were sonicated for 25 cycles of 30 seconds on and 30 seconds off. A 30 µL aliquot was crosslinked to be used as input. The remaining mixture was centrifuged at 4°C for 15 minutes, and the supernatant was transferred into a new tube and frozen at −80°C. The immunoprecipitation was done by adding antibody (2 ug of H3K4me3 antibody, Active Motif, #39159), then rotated overnight at 4°C. Next, 40 µL of beads (DynabeadsTM Protein A, Invitrogen, REF #10004D) were taken and washed twice with 1 mL of FA + PI. The beads were resuspended in 1 mL of FA + PI + 1% BSA + 10 µL of tRNA and rotated overnight at 4°C. The beads were washed twice with FA + PI and then transferred into the extract/antibody solution, followed by rotation at 4°C for 2 hours. The beads were then washed twice in FA + PI, once in FA with 500 mM NaCl, once in FA with 1 M NaCl, and twice in TEL buffer (0.25 M LiCl, 1% IGEPAL, 1% sodium deoxycholate, 1 mM EDTA, 10 mM Tris-HCl pH 8). The beads were eluted in 60 µL of ChIP elute buffer and incubated at 65°C for 15 minutes. The elution was transferred to a new tube as the IP. For decrosslinking, 2 µL of RNase (Roche, 1119915001) was added, and the mixture was incubated at 37°C for 1.5 hours. Finally, 1.5 µL of proteinase K (20 mg/mL, NEB, P8107S) was added, and the mixture was incubated overnight at 65°C for decrosslinking. The DNA was purified from the solution using a PCR purification kit (Invitrogen, K31002).

DNA libraries were prepared and sequenced by Novogene. The purified DNA samples were treated with End Repair Mix (Novogene) and incubated at room temperature for 30 minutes. They were then purified using a PCR purification kit (Qiagen). The DNA was subsequently incubated with A-tailing mix at 37°C for 30 minutes. Next, the 3’-end adenylated DNA was ligated with the adapter in the ligation mix at 20°C for 15 minutes. The adapter-ligated DNA underwent several rounds of PCR amplification and was purified using a 2% agarose gel to recover the target fragments. The average fragment length was assessed using an Agilent 2100 Bioanalyzer (Agilent DNA 1000 Reagents) and quantified by qPCR (TaqMan probe). The libraries were further amplified on a cBot system to generate clusters on the flow cell and sequenced on an Illumina Novaseq X plus system (paired-end 50 base read length).

Adapter sequences were removed using Cutadapt 1.18^108^ in the pair-ended read mode. Reads were then mapped to the *C. elegans* genome (WBcel235) using the Burrows-Wheeler Aligner with the MEM algorithm (BWA 0.7.17-r1188)^109^. Mapped reads were indexed and sorted using Samtools 1.10^110^. Bam files were filtered with samtools to remove non-unique mappers, secondary alignments and low-quality pairs (MAPQ<10). Duplicate reads were removed using Picard MarkDuplicates 3.1.0-3^111^ with the --REMOVE_DUPLICATES option. Peak calling was performed using Macs3 CallPeak (v3.0.0b1)^112^ with no cutoff (-q 1) and the options --extsize 200 and --nomodel. Differential binding analysis was conducted using DiffBind (v3.12.0)^113^ in R (v4.3.3) over a 200 bp sliding window with 10 bp shift across the entire genome. Counts were obtained without merging overlapping peaks and without computing summits. Normalization was performed using the DESeq2 method^114^, accounting for library size, and differential analysis followed DiffBind’s DESeq2 implementation.

BedGraph files displaying enrichment normalized to input for wild-type and *set-24* mutants, along with fold-change between *set-24* and wild-type enrichment, were generated using a custom Python script with a bin size of 10 bp. Metagene analysis was performed over all protein-coding genes identified in Ensembl’s annotation (release 112) for WBcel235. The average H3K4me3 enrichment at TSS was calculated within a ±500 bp window. Genes contributing to 80% of the total enrichment were selected for further analysis and classified as H3K4me3-enriched genes. All figures were generated using custom Python scripts. Analyses were performed using Python (v3.8.10) with Pandas (v2.0.1)^115^ for data management, NumPy (v1.23.5)^116^ for calculations, SciPy (v1.3.3)^117^ for statistical tests, and Matplotlib (v3.4.3)^118^ for figure generation.

### mRNA-seq and sequencing analysis

mRNAs were purified from total RNA using PolyT oligo-attached beads and converted to cDNAs for library preparation. cDNA libraries were sequenced using a paired-end 150 bp sequencing strategy on an Illumina Novaseq X plus system. Raw reads were assessed for quality using FastQC, Picard Tools, and Samtools along with trimming of poor-quality reads using Trimmomatic (v0.39; paramters: SLIDINGWINDOW:4:20, MINLEN:36, ILLUMINACLIP:TruSeq2-PE.fa:2:30:10)^110,119,120^. Clean reads were processed to find transcript abundance counts with Salmon (v1.10.2; paramters: --gcBias, --seqBias)^121^. We used DESeq2 (v1.46.0; padj < 0.01) to identify differentially expressed genes in *set-24* mutants compared to the WT^114^.

### small RNA-seq and sequencing analysis

Total RNAs were treated with RppH (NEB #M0356S)^122^, and small RNA libraries were prepared with a small RNA-seq Kit v4 with UDIs (Nextflex #NOVA-5132-31). Libraries were sequenced using a single-end 50 bp sequencing strategy on a Novaseq6000 platform. Adapters and reads shorter than 18 nucleotides were removed using CutAdapt v1.15^123^, with options -a TGGAATTCTCGGGTGCCAAGG --minimum-length 18. The quality of raw and trimmed reads was assessed with fastQC v0.11.9^119^. Subsequently, 22G-RNAs, defined as all reads between 21 and 23 nucleotides long starting with a G, were isolated. This filtering was conducted using a combination of CutAdapt v1.15^123^, with options --minimum-length 21 --maximum-length 23, and zcat/awk utilities. 22G-RNAs were mapped to the *C. elegans* genome (WBcel235) using STAR v2.7.3a^124^, with options --readFilesCommand zcat --outMultimapperOrder Random -- outFilterMultimapNma× 100 --outFilterMismatchNmax 0 --alignIntronMax 1 --outSAMtype BAM SortedByCoordinate --outFilterType BySJout --winAnchorMultimapNma× 100 -- alignEndsType EndToEnd --scoreDelOpen −10000 --scoreInsOpen −10000 –outSAMmultNmax 1. Subsequent quantification of counts and differential expression analysis were conducted as previously reported^125^. In short, featureCounts v2.0.0^126^ was used to calculate counts mapping to genes (-t exon), using the BAM files produced in the previous STAR alignment step as input. The resulting tables of counts were imported into R, and DESeq2^114^ and custom scripts (available online^127^) were used to obtain normalised counts and conduct statistical tests. The following R packages were used: tidyverse^128^, lattice^129^, eulerr^130^, genefilter^131^, reshape2^132^, ashr^133^, GenomicFeatures^134^. Genes with differentially expressed 22G-RNA levels in *set-24(mj617)* mutants, defined by fold change > 2 and 5% FDR, were overlapped with known target genes of specific small RNA populations using BioVenn^135^. We used previously described lists of known target genes^136–140^.

### Generation of genome tracks

bedGraph files with ChIP-sequencing read counts normalised to library size and to input were converted to bigWig format using bedGraphToBigWig v2.8^141^. Small RNA and mRNA bigwig files and genome tracks were created as previously described^125^. In short, bigWigs were generated with bamCoverage v3.5.1^142^, using options --normalizeUsing CPM --binSize 5 (for small RNA) or –binSize 10 (for mRNA) and the BAM files with 22G-RNAs/mRNAs mapped to the *C. elegans* genome. All the replicates of the same strain, either wild-type N2, *set-24(syb7014)*, or set-24(*mj617*) mutants, were combined using WiggleTools mean^143^ and wigToBigWig v4^141^. Genome tracks were plotted with custom scripts^127^, on an R framework (R Core Team 2021), using the Gviz^144^ and GenomicFeatures^134^ packages.

### Data sets

Sequencing data have been deposited to the NCBI Gene Expression Omnibus (GEO), and proteomics data are available at the ProteomeXchange Consortium via PRIDE. GEO: GSE291568, PRIDE: PXD057349.

### Statistics

All experiments were conducted with independent *C. elegans* animals for the indicated N times. Statistical analysis was performed as indicated in the figure legends.

## Acknowledgements

We sincerely thank all members of the Miska lab for their valuable feedback on this work. We are grateful to the *Caenorhabditis* Genetics Center (CGC), funded by the National Institutes of Health Office of Research Infrastructure Programs (P40 OD010440), and the National Bioresource Project (NBRP) for providing the strains used in this research. We also extend our appreciation to Marie-Anne Félix’s and Siu Sylvia Lee’s labs for sharing worm strains and unpublished data. Special thanks go to Helena Santos-Rosa Ruano for her assistance with the detailed analysis of the SET-24 sequence. We are also thankful to Yuhua Lim and Lisa Lampersberger for their comments and support. This research was supported by grants from Wellcome Trust Senior Investigator Award (219475/Z/19/Z) and CRUK award (C13474/A27826). We also acknowledge core funding to the Gurdon Institute from Wellcome (092096/Z/10/Z, 203144/Z/16/Z) and CRUK (C6946/A24843).

## Author contributions

G.F. and E.A.M. conceived the study. C.Z., G.F., M.V.A., J.C.R., and J.M. conducted the experiments. J.P., J.C.R., and M.V.A. analysed the sequencing data. M.V.A and P.R.-G. performed evolutionary and structural analysis of SET and SPK domains. M.H. performed the western blotting under the supervision of S.G. F.B. processed samples and conducted mass spectrometry. E.A.M. supervised and funded the study. C.Z. drafted the initial manuscript with contributions from coauthors, and all authors reviewed and edited the final version.

---

**Figure S1. Related to Figure 1.**
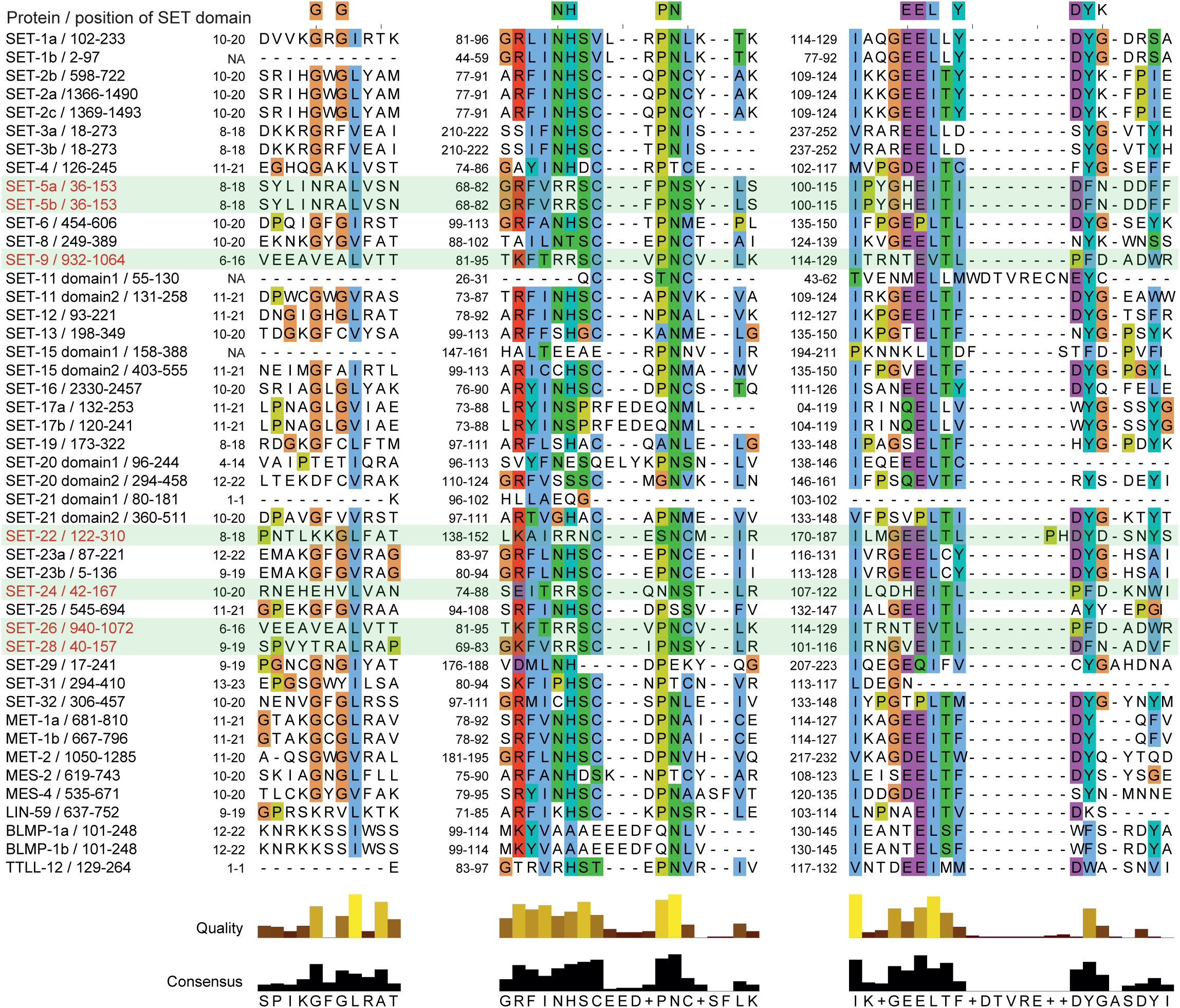
Alignment of *C. elegans* SET domains. Multiple sequence alignment of SET domains. Residues critical for catalytic activity are highlighted. SET proteins in the branch highlighted in Fig. 1c are indicated with red labels and green background.

**Figure S2. Related to Figure 1.**
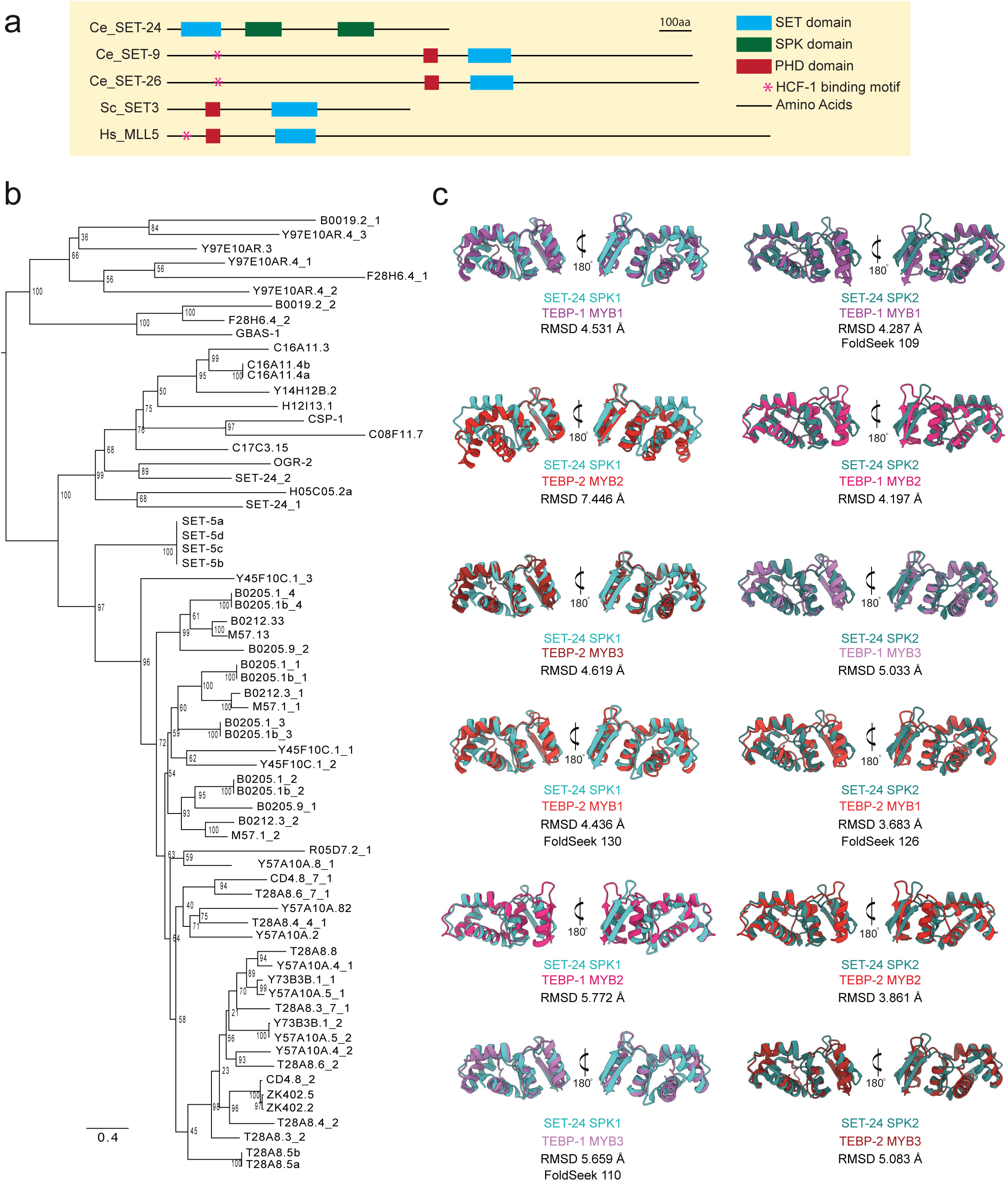
Evolutionary and structural analysis of SPK domains. a Schematic representation of sequences of SET-24 and Set3 subfamily proteins with domain information. b Maximum likelihood phylogenetic tree comparing the protein sequences of SPK domains in *C. elegans*. Values next to the tree nodes are branch supports, calculated with 1,000 ultrafast bootstrap replicates. c The structural alignments show the similarity between the SPK domain of SET-24 and the MYB domains of TEBP-1 and TEBP-2, as predicted by Alphafold3.

**Figure S3. Related to Figure 2.**
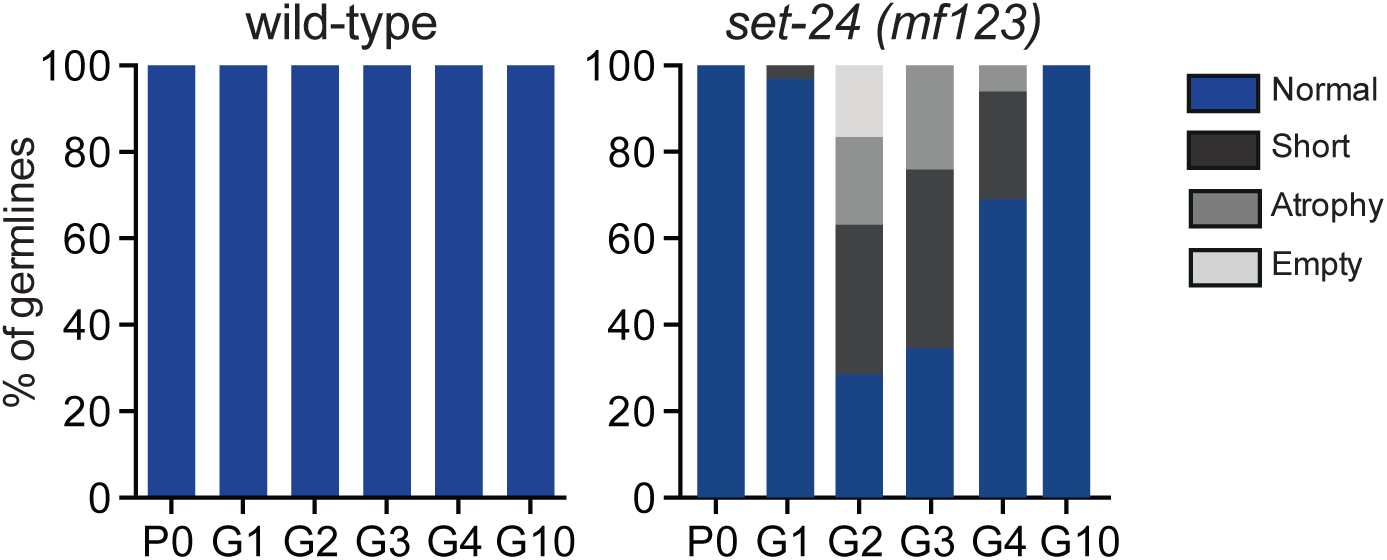
SET-24 is required for germline development. *set-24(mf123)* mutant worms, the same mutation isolated from the wild strain display progressive germline degeneration. Proportions of normal, short, atrophic, and empty germlines (countings on pooled progeny of 15 parents. n>30 per genotype).

**Figure S4. Related to Figure 3.**
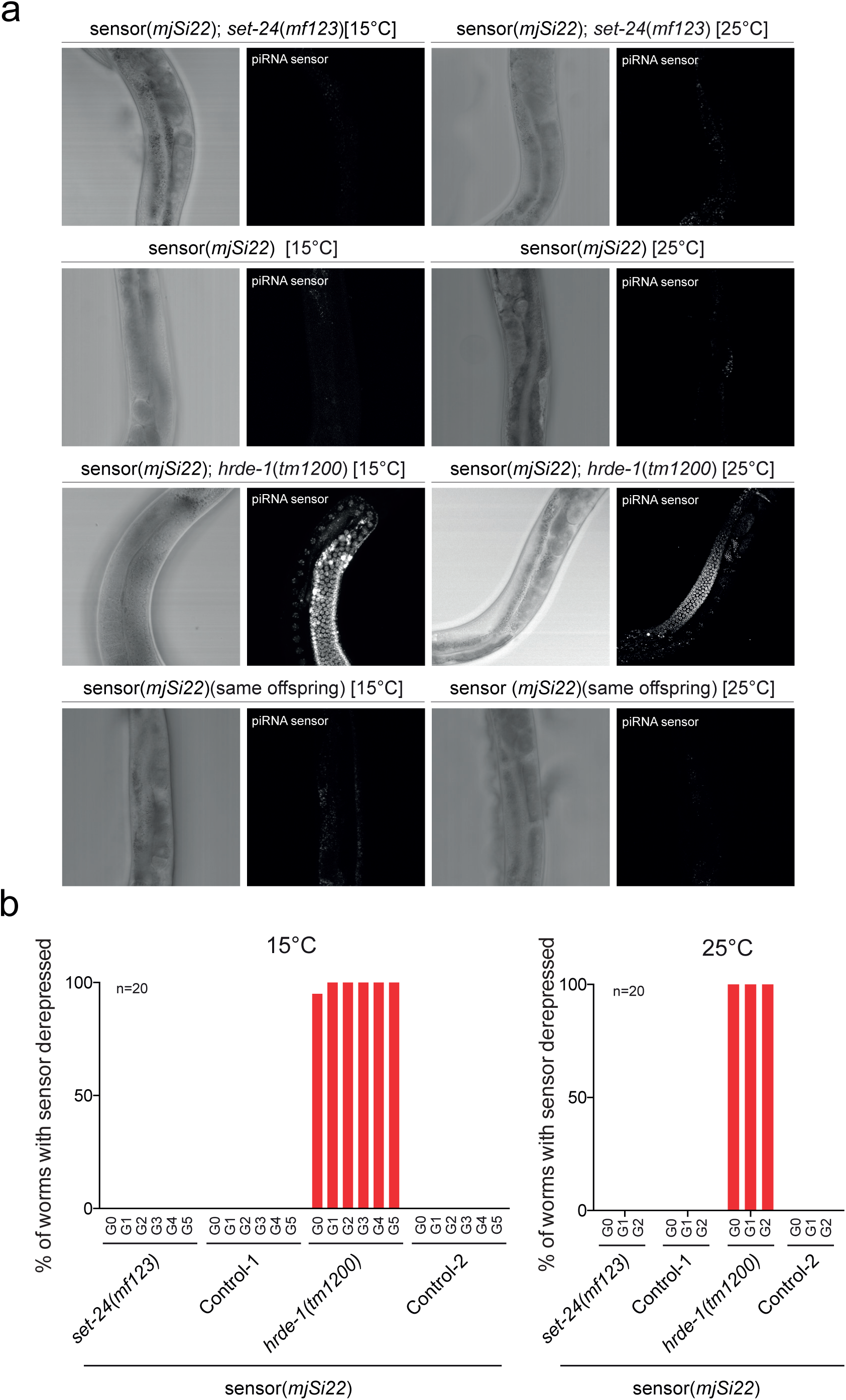
*set-24* is dispensable for initiation of piRNA-dependent silencing. **a** Representative images of G2 generation germlines of the indicated genotypes. **b** Quantification of the percentage of derepressed individuals at every generation (countings on n > 35 animals per generation).

**Figure S5. Related to Figure 4.**
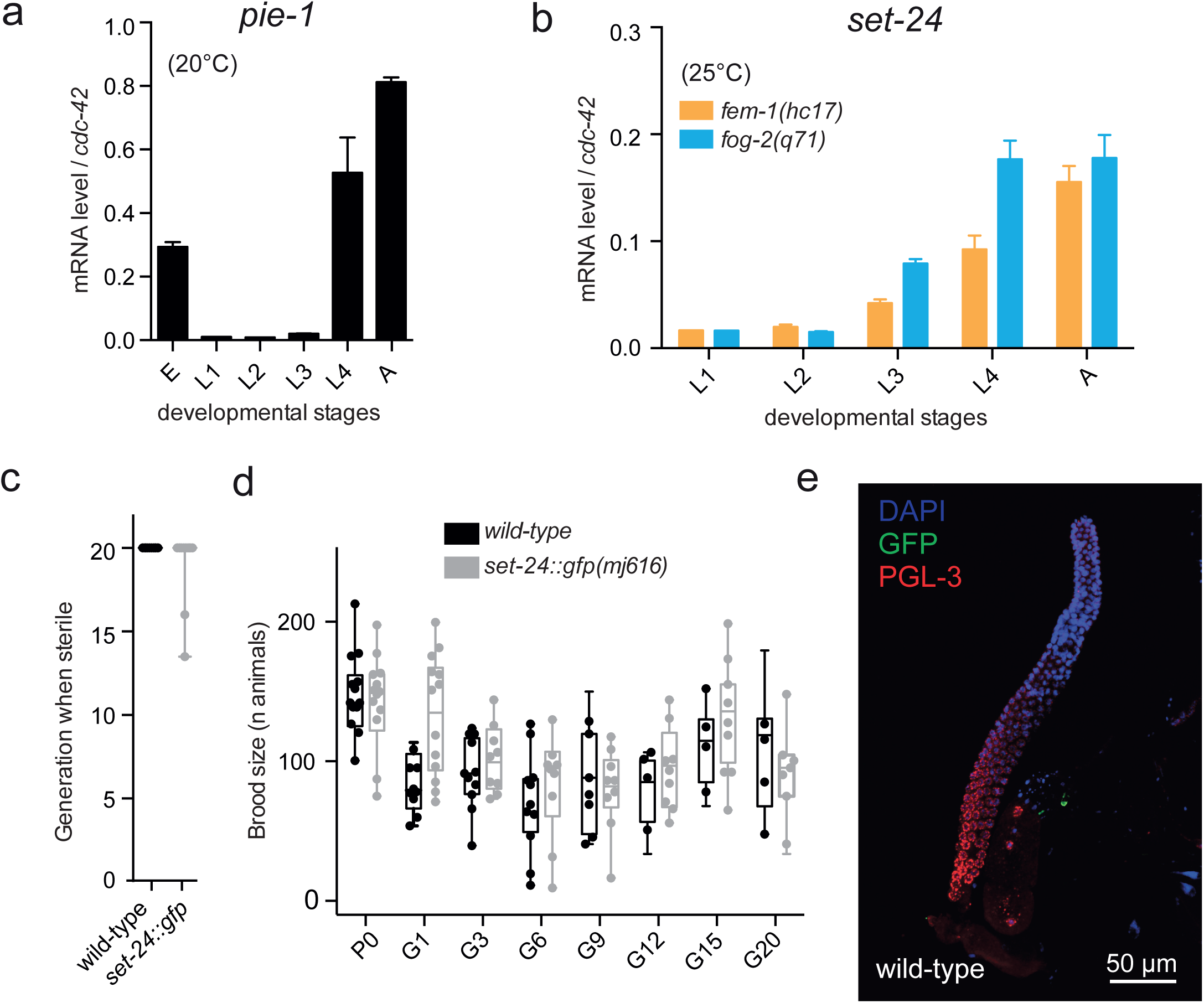
SET-24 is a germline-specific factor. **a** and **b** RT-qPCRs showing means and standard deviations over three independent experiments. **a** Developmental time-course of *pie-1* expression in *wild-type* animals grown at 20°C. **b** Expression analysis of *set-24* in masculinized (*fem-1(hc17)*) and feminized (*fog-2(q71)*) mutants grown at 20°C. **c** and **d** SET-24::GFP animals are not germline mortal at 25°C. Assessment of fertility, n=15 (**c**) and quantification of the number of progeny over generations (P0, parental generation; G1-G20, generations 1 to 20) n=15 per genotype (**d**). **e** Representative immunofluorescence image of control wild-type animals stained with anti-GFP and anti-PGL-3 antibodies. Compare with Figure 4c.

**Figure S6. Related to Figure 5.**
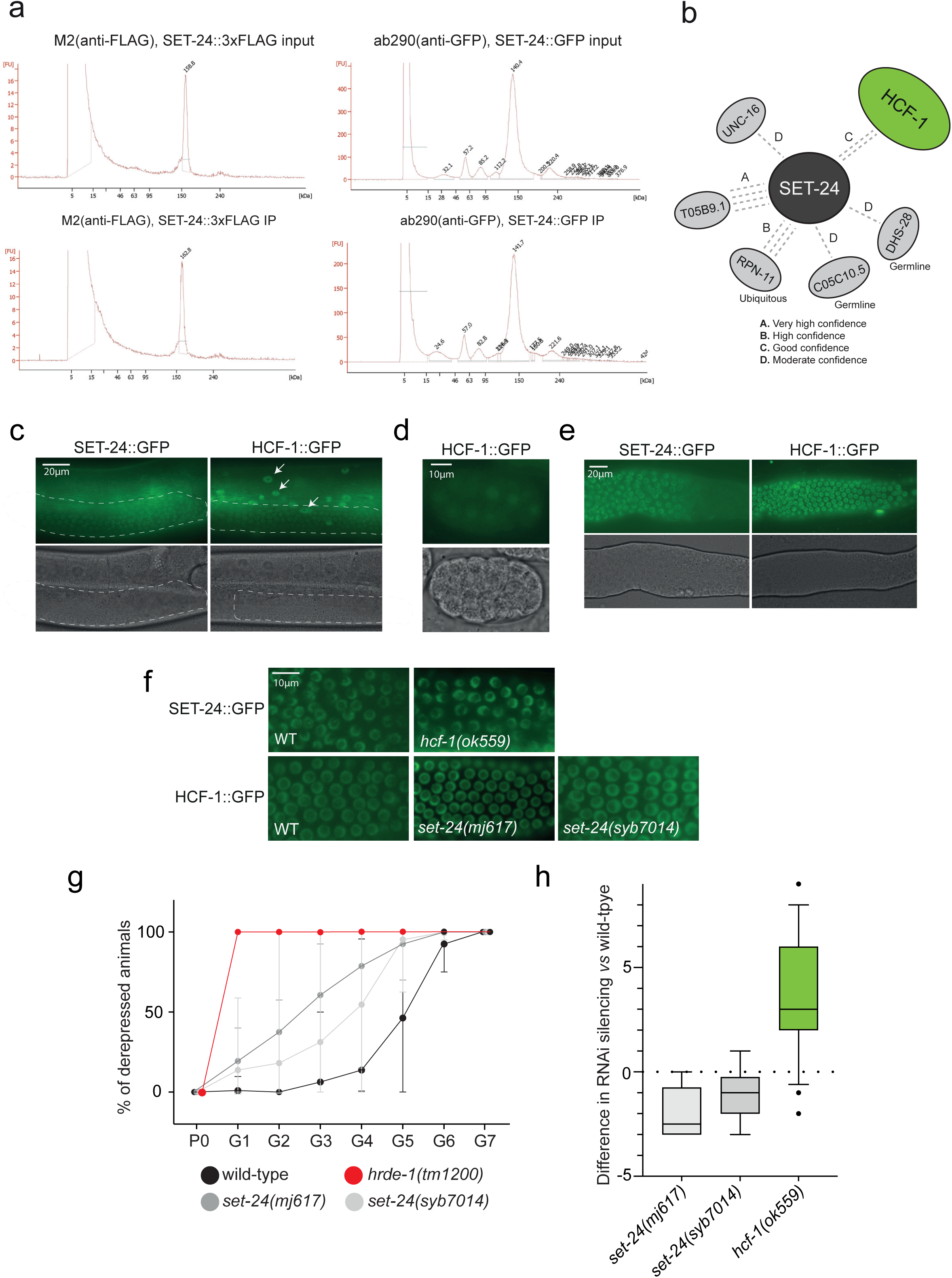
The expression of SET-24 and HCF-1. **a** Profiles of DNA from SET-24::3xFLAG or SET-24::GFP ChIP and input, analysed using TapeStation. **b** Identification of SET-24 interactors via Yeast-two Hybrid. Panels “A-D” indicate the confidence levels of interaction. **c** Representative images showing somatic expression of GFP-tagged SET-24 and HCF-1, marked by white narrows. The germline marked by a white dashed line. **d** A representative image showing the expression of GFP-tagged HCF-1 in the embryo. **e** Representative images showing expression of GFP-tagged SET-24 and HCF-1 in the extruded gonads. **f** Representative images of germlines of SET-24::GFP and HCF-1::GFP in wild-type and *hcf-1* or *set-24* mutant background. **g** Quantification of the percentage of derepressed individuals at every generation. n > 6. **h** The contrasts between generations of derepressed mutants and their corresponding wildtypes. The y-axis represents the difference in the number of generations of silencing between mutants and wildtypes. For instance, if silencing (defined as more than 50% of worms remaining silenced) continues until G8 in mutants but only until G5 in wildtypes, the contrast would be 3. *set-24* mutants, n > 6; *hcf-1(ok559),* n > 20.

**Figure S7. Related to Figure 6.**
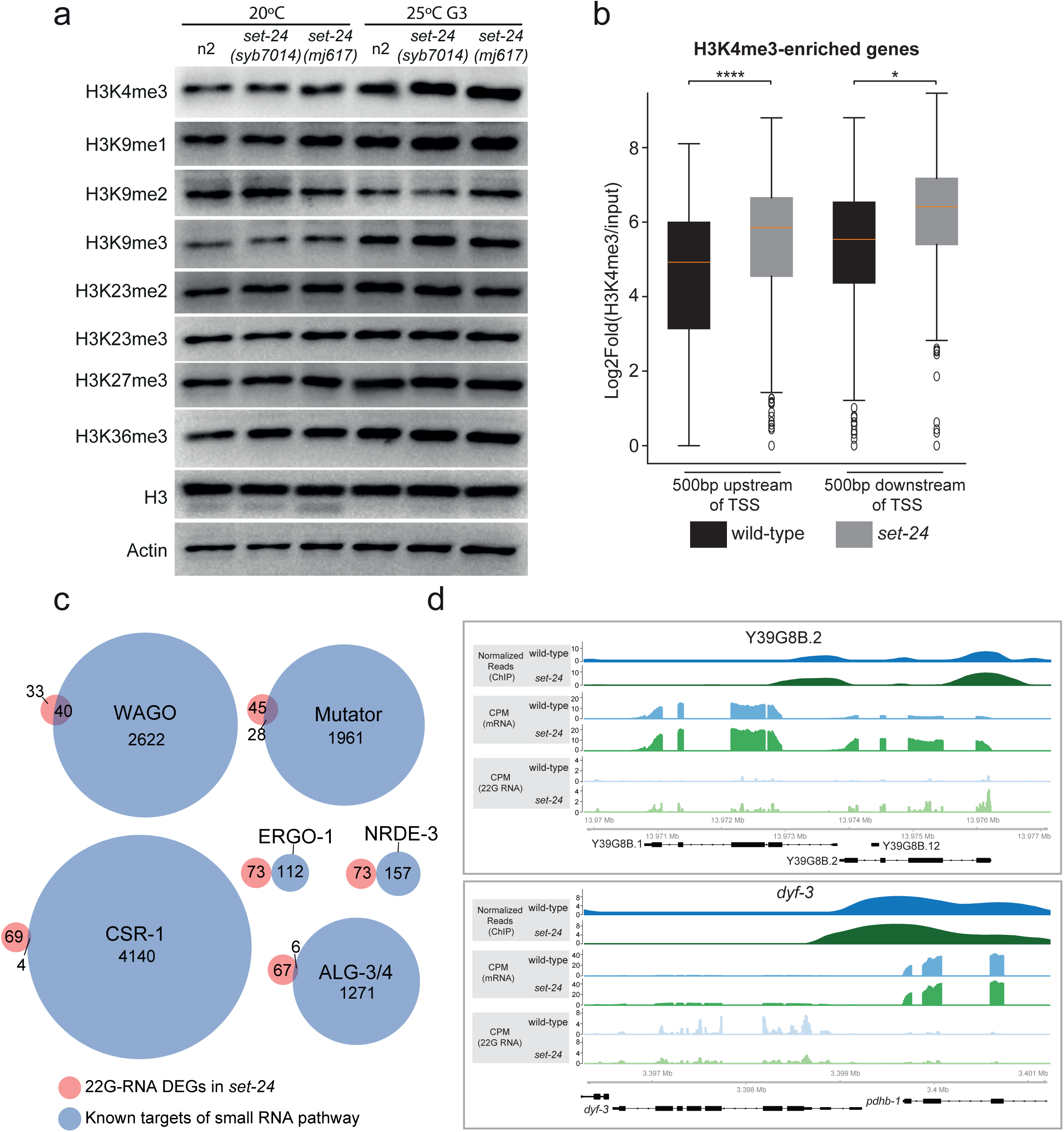
SET-24 is required to regulate H3K4me3 and small RNA levels. **a** Western blot analysis of wild-type and *set-24* mutant worms was performed using the specified antibodies. Worms were collected at the young adult stage, either grown at 20°C or after being cultured at 25°C for three generations. **b** Comparison of average H3K4me3 enrichment within 500 bp upstream and downstream of TSS for wild-type samples and *set-24(syb7014)* mutants across H3K4me3-enriched genes with significant log₂ fold-change. Statistical significance was assessed using a paired *t*-test. Asterisks indicate p-values: ****p < 0.001; *p < 0.05. **c** Venn diagrams showing overlaps between lists of genes with deregulated 22G RNA levels in in *set-24(mj617)* mutants (fold change > 2 and 5% FDR) and genes known to be targeted by specific small RNA pathways. **d** H3K4me3 enrichment, mRNA, and 22G RNA levels of the selected genes (Y39G8B.2 and *dyf-3*) in wild-type and *set-24* mutant strains.

**Supplementary Table 1**. Immunoprecipitation-Mass spectrometry results of anti-FLAG IPs on N2 (expN) and SET-24::3xFLAG (expS) young adult worms.

**Supplementary Table 2.** Yeast Two Hybrid screening results of SET-24 protein.

**Supplementary Table 3.** Log2fold change of mRNAs in *set-24(syb7014)* compared to wild-type.

**Supplementary Table 4.** Lists of H3K4me3 enriched genes with upregulated mRNAs, upregulated 22G-RNA targets or downregulated 22G-RNA targets.

**Supplementary Table 5.** WormExp enrichment analysis of H3K4me3 enriched genes with upregulated mRNAs, upregulated 22G-RNA targets, or downregulated 22G-RNA targets.

**Supplementary Table 6.** Log2fold change of 22G-RNAs in *set-24(mj617)* compared to wild-type.

**Supplementary Table 7.** Worm strains used in this work

